# Directional uncertainty in chase and escape dynamics

**DOI:** 10.1101/2023.08.06.552194

**Authors:** Mario Treviño, Ricardo Medina-Coss y León, Sergio Támez, Beatriz Beltrán-Navarro, Jahir Verdugo

## Abstract

Intercepting and avoiding collisions with moving targets are crucial skills for survival. However, little is known about how these behaviors are implemented when the trajectory of the moving target introduces variability and ambiguity into the perceptual-motor system. We developed a simple visuomotor task in which participants used a joystick to interact with a computer-controlled dot that moved along two-dimensional trajectories. This virtual system allowed us to define the role of the moving object (predator or prey) and adjust its speed and directional uncertainty (*i.e.,* magnitude and frequency of random directional changes) during chase and escape trials. These factors had a significant impact on participants’ performance in both chasing and escaping trials. We developed a simple geometrical model of potential chaser/escaper interactions to distinguish pursuit from interception chasing trajectories. We found that participants initially pursued the target but switched to a late interception strategy. The amount of late interception strategy followed an inverted U-shaped curve with the highest values at intermediate speeds. We tested the applicability of our task and methods in children who showed a robust developmental improvement in task performance and late interception strategy. Our task constitutes a flexible system in a virtual space for studying chasing and escaping behavior in adults and children. Our analytical methods allow detecting subtle changes in interception strategies, a valuable tool for studying the maturation of predictive and prospective systems, with a high potential to contribute to cognitive and developmental research.

## Introduction

Animals use a broad set of maneuvers to chase moving prey and escape from predators. Sensory and locomotor performance influence the outcome of these confrontations (Domenici et al., 2011; Ghose et al., 2006). At least two main chasing approaches can be identified: pursuit and interception (Mischiati et al., 2015). Pursuing refers to actively following a moving target, which entails keeping it constantly in the same field of view while advancing towards it in the same direction. Interception, in contrast, is a more complex approach since it involves predicting the target’s future location while advancing toward that predicted location. Real-world object interception requires deciding when to start and direct a movement based on a forecast of motion sources (*i.e.*, those objects that initiate or generate motion) and then making continuous adjustments due to changing conditions in the outside world. Thus, several sources of information and coordination across multiple sensory-motor systems are necessary to predict the kinetics of interception. Not surprisingly, predators that intercept well are less likely to rely on an absolute speed advantage, which helps them save energy (Festa-Bianchet & Mysterud, 2018; Mischiati et al., 2015).

The ability to intercept moving targets and prevent collisions is also essential for daily survival in humans. Many activities require localizing and directing movements toward a stimulus, using perceptual inputs such as touch, vision, and proprioception (Chinn et al., 2019). Games like tennis and baseball require precise visual information about when to intercept a ball based on its direction and velocity (Brenner & Smeets, 2015). Other day-to-day activities, such as car driving, depend upon good movement predictions to avoid collisions. All these activities impose severe spatial-temporal constraints that push biological systems to develop robust and flexible perceptual-motor behaviors (Franklin & Wolpert, 2011; Ghose et al., 2006; Mischiati et al., 2015; Niehorster et al., 2015; Soechting et al., 2009; Wolpert et al., 1995).

Numerous studies have investigated visuomotor coordination with stationary and moving targets (Fooken et al., 2021; Mrotek & Soechting, 2007; Zago et al., 2009). Target speed directly influences interception and evasion actions. Directional uncertainty in the moving target adds complexity by introducing variability and ambiguity into the perceptual-motor system. However, the specific adaptations that occur in tasks with varying levels of directional uncertainty during chasing and escaping conditions have not been thoroughly investigated. We hypothesized that the targets’ speed and degree of directional uncertainty should lead to distinct behavioral patterns in movement trajectories and collision times.

To address this question, we developed a novel computer-based task that allowed us to capture participants’ trajectories and kinematics using a joystick, enabling us to examine the overall functionality of the visuomotor system interacting with a moving object. The task involved chasing or escaping from a computer-controlled moving target displayed on a 2D screen (*i.e.,* the task did not involve depth perception). Participants observed the dot’s movement and interacted with the target from a third-person perspective. We aimed to develop flexible and reliable conditions for studying directed visuomotor behaviors. While our task enables the exploration of different chasing/escaping approaches, this initial investigation serves as a proof-of-concept, demonstrating the capability of our system to capture and analyze visuomotor behaviors under controlled uncertainty. We had the flexibility to define the role of the moving object (predator or prey) and adjust its speed while manipulating the magnitude and frequency of random directional changes, which introduced varying degrees of uncertainty in the target’s path. We thus characterized how these variables determined chasing (collision achievement) and escaping (collision avoidance) performance. We found that decreasing target speed resulted in improved chasing and escaping performance, while directional uncertainty had a negative impact on chasing but a positive impact on escaping performance.

We developed a methodology to analyze chasing trajectories differentiating pursuit from interception strategies. Applying this approach, we found that skilled chasers showed a greater tendency for a late interception strategy, which became more prevalent as the uncertainty in the future target motion/location decreased, particularly near the collision point. In contrast, less proficient chasers exhibited a lower usage of interception, leading to reduced success rates during attacks. Additionally, we explored the task’s suitability in younger participants, revealing a significant developmental increase in task performance and utilizing the late interception strategy across 6, 10, and 20-year-old groups.

We explored how individuals deal with directional uncertainty when chasing or escaping from moving targets. We developed an innovative approach to investigate visuomotor strategies and demonstrate their feasibility and potential. Initially, our experiments served as a proof-of-concept, highlighting the capabilities and possibilities of the approach for studying visuomotor coordination. Our task and methods can be effectively utilized to explore visuomotor behaviors across diverse populations, providing clinicians and scientists with a valuable tool to assess the underlying processes of chasing and escaping behavior. Additionally, our approach holds promise in differentiating between neurotypical and neurodivergent adults and identifying potential disabilities in children.

### Public Significance Statement

We developed a computer-based visuomotor task to investigate how people chase and escape from moving targets under different conditions. Lower target speeds improved chasing and escaping abilities, while higher directional uncertainty compromised chasing but improved escaping. This research enhances our understanding of how humans adapt their behavior to intercept or avoid moving objects. Our approach has cognitive and developmental research applications, as children showed developmental improvements in task performance and strategy utilization. For example, clinicians could use a similar methodology to assess visuomotor performance and identify individual impairments. It has the potential to differentiate between neurotypical and neurodivergent adults and identify developmental disabilities in children, aiding in diagnostic tools and interventions for visuomotor difficulties. By capturing these behaviors’ complexity and dependence on speed and uncertainty, our research contributes to our understanding of visuomotor coordination in children and adults.

## Method

### Transparency and openness

We are committed to ensuring transparency and openness in our research. We make raw data and supplementary materials available through the Open Science Framework (OSF, https://osf.io/nsbhr) to facilitate replication and verification. Detailed descriptions of our experimental procedures, data collection protocols, and data processing steps are provided in the following sections to ensure methodological transparency. Participant data was treated as confidential, following legal and ethical guidelines, and used solely for research purposes. Participants and/or their guardians, in the case of minors, were informed about the research findings upon request. We respected participants’ rights to be informed about the study’s outcomes. We have justified our sample size and reported all manipulations and measures. We excluded nine outlier participants from the data analysis (see below). We disclose any potential conflicts of interest that may impact the interpretation or outcomes of our study at the end of the manuscript. All studies conducted in this research were approved by our University (see below).

### Participants

We conducted experiments on 295 adults (157 women, 138 men; age range (yr.): 19 - 30, mode: 22), and 40 children/youngsters with ages between 6 and 10 years old (21 girls, 19 boys). We formed three age groups (6, 10, and 20) based on the developmental patterns of gray matter growth in the human brain. Namely, between the ages 4 and 11, gray matter exhibits a relatively linear developmental course, yet, by the age of 20, the total brain size reaches 95% of its maximum size, indicating a period of greater stability. Thus, by focusing on these specific age groups, we aimed to capture behavioral changes potentially associated with critical gray matter development stages during childhood and early adulthood (Lenroot & Giedd, 2006). We used guidelines of typical development as inclusion criteria: i) born at term, ii) birth weight between 2500 and 3999 grams (5.5-8.8 lbs.), iii) no reports of prenatal, perinatal, or postnatal complications or trauma that could affect nervous system development. No participant was diagnosed with a psychiatric, neurological, or neurodevelopmental disorder or had a history of previous or current substance use. The inclusion criteria and clinical questionnaires were used to control their ‘typical’ status. Most participants were dextral (≥ 98 %, *i.e.,* corresponding to their writing hand) and had a normal or corrected-to-normal vision (Snellen test; (Trevino et al., 2020)). We gave adult participants written instructions on responding to the questionnaires and solving the task. For children, the questionnaires were answered by their parents. These formularies involved demographic and clinical history questions. Participation was strictly voluntary and involved no monetary incentives. This work was performed in accordance with all local guidelines and regulations of our state, and all the methods were approved by the ethics committee of Instituto de Neurociencias: ET102021-330, and Centro Universitario de Ciencias de la Salud, CUCS: CI-04022; Universidad de Guadalajara, México.

### Collection of demographic information from participants

Our project adhered to fundamental principles of respect for individuals, beneficence, non-maleficence, and justice. It followed international ethical standards for biomedical research involving human subjects. We considered the regulations outlined in various articles of the General Health Law for Research in Mexico, the Helsinki Declaration, and the guidelines established by the Council for International Organizations of Medical Sciences (CIOMS). The study also adhered to the criteria for executing research projects involving human subjects outlined in the NOM-012-SSA3-2012. To ensure ethical compliance, we implemented measures to protect participants’ data confidentiality, conducted risk identification and mitigation and obtained informed consent and assent. All adult participants provided written informed consent. For minors, we obtained parental consent and explicit assent from the children. These steps were taken to uphold ethical standards and safeguard the rights and well-being of the participants. We got written permission to take and ‘cartoonize’ the pictures from participants illustrated in *Figure 1a*, and *Figure 6a*. To ensure participant well-being, breaks were provided, and data confidentiality was strictly maintained. Participants could withdraw from the study at any moment without penalty and request the deletion of their data. The questionnaires utilized open-ended response formats, allowing participants to provide detailed answers. No questions regarding race or ethnicity were included. The testing was conducted individually in controlled environments with standard lighting and without distractors. Participants had a comfortable seat height to view the screen, and both participants and experimenters maintained a quiet environment. Sessions took place in designated spaces provided by the institutions.

**Figure 1.**
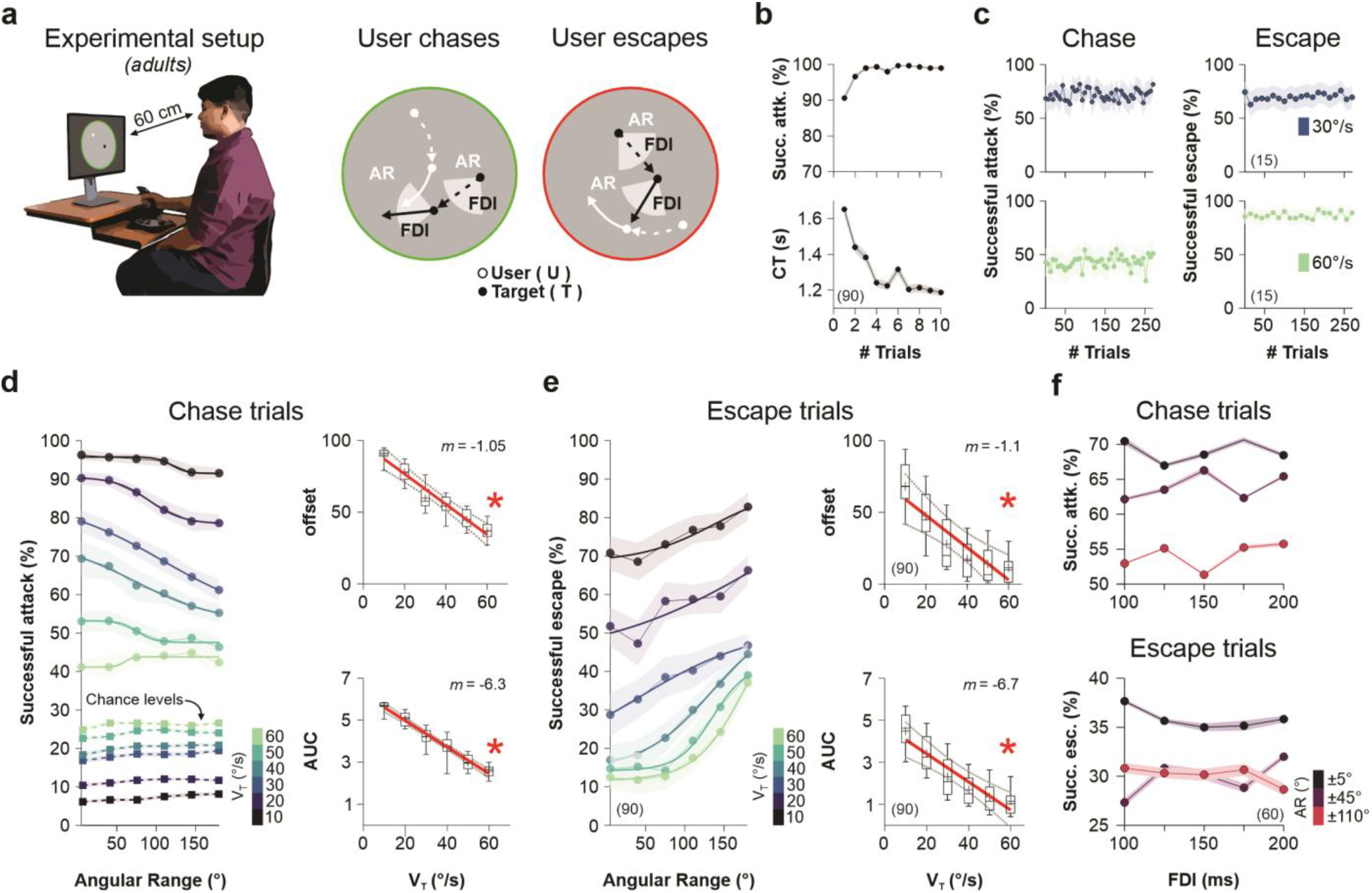
Minimalistic visuomotor task to study chasing and escaping behavior. **a** Cartoon of the experimental setup. Each participant sat in a chair and controlled the movement of a white dot using a joystick. Participants could adopt one of two roles: chase the target or escape from it. The target, displayed as a black dot, moved in rectilinear trajectories, changing direction every ‘fixed-direction interval’ (FDI). The change in direction was determined using random numbers from a uniform distribution with a user-controlled angular range (AR). **b** Participants were pre-trained to ensure that were able to properly use the joystick. **c** Stable chasing and escaping performance during the entire experimental session. **d,e** Group psychometric curves for successful attacks (left) and escapes (right) performed with an FDI = 100 ms. Target speeds (*v_T_*) depicted in the colorbar. Offsets and area under the curve (AUC) of the sigmoids as a function of *v_T_* are shown on the right side of each panel. Chance levels depicted as squares with dotted lines in panel **d** (chance levels for successful escapes are shown in *Supplementary Figure 1*). Asterisks depict significant slopes (*P* ≤ 0.05). **f** Chasing (top) and escaping (bottom) performance for different FDIs tested. Colors correspond to different AR groups depicted on the colorbar. Number of participants per experiment in parentheses. Asterisks depict significant slopes (*P* ≤ 0.05).

### Visuomotor task

Participants sat unconstrained in front of a computer to solve a visuomotor task involving chasing or escaping from a computer-controlled object. The visual stimuli were two small moving dots of 0.3° of visual angle: a white dot representing the participant (○ controlled by moving a joystick with the dominant hand) and a black dot representing the computer-controlled object (●). Both dots were displayed simultaneously within a circular arena (radius ≈ 14.82 cm ≈ 540 pixels; 50% gray background) placed on the center of a single 27-inch computer monitor (Dell P2414H; at a viewing distance of ∼60 cm; *Figure 1a*, *Figure 6a*). The monitor had a resolution of 1920 × 1080 pixels (∼52.70 cm × ∼29.64 cm) and operated at a 60 Hz refresh rate, leading to an acquisition speed of one frame every ∼16.66 ms. We recorded the responses of the participants using a joystick (Logitech Extreme Pro 3D, 1000 Hz (Trevino et al., 2020)) connected to a regular computer (Intel (R) Xeon (R) @ 3.40 GHz; 64-bit operating system; NVIDIA Quadro K600, 8 G.B. graphics card). The joystick was calibrated using a five-step routine in the Windows control panel, and a MATLAB R2022 (MathWorks, Inc.; Natick, USA) program read out its relative position. The joystick gain was set to one, ensuring a linear relationship between the joystick inclination and the pixel displacement of the white dot on the screen, with the joystick’s position being able to reach the edges of the screen. At the beginning of every trial, the participant (○) appeared at the center of the screen. In contrast, the computer-controlled target (●) appeared in a random position along the circumference of the arena (uniform distribution to generate randomly permuted initial angles). Subsequently, the target moved in sequences of rectilinear trajectories (see below) within the arena, corresponding to the area delimited by the circumference. Depending on visual instructions during each trial, the participant could either chase or escape from the target (throughout the manuscript, we will refer to the black dot controlled by the computer as the ‘target’). Thus, ‘chase trials’ were those in which the participant had to chase the moving target (○→●), whereas ‘escape trials’ involved escaping from it (*i.e.,* avoiding being hit by the black dot: ●→○). These two conditions were randomly permuted and shown at the beginning of each trial by using either a green (chase trials) or a red (escape trials) colored ring around the arena (right panel on *Figure 1a*). The task instructions emphasized achieving successful attacks/escapes (rather than speed). Trials terminated when both dots touched each other externally (*i.e.,* spatial and temporal coincidence of the dots) or when the trial duration exceeded 5 s. A ‘successful attack’ refers to a chase trial in which the participant effectively collided with the target, whereas a ‘successful escape’ involves an escape trial in which the participant avoided being hit by the target for 5 s. We included auditory feed-back using tones with pure frequencies to indicate successful (10 kHz) and unsuccessful (1 kHz) trials. We measured the collision time (CT, in s) as the interval between the appearance of the dots at the beginning of the trial and their collision (CT = 5 s for successful escape trials). Thus, task efficiency was associated with shorter CTs in chase trials but longer CTs in escape trials. As an exclusion criterion, we removed data from 9 outlier participants who showed ≥ ±10% change in average CT during the resolution of the task. Some of these participants displayed anomalous behaviors that were not explicitly instructed (such as lifting the joystick into the air and forcefully striking it against the table). After finishing a trial, the participants had to restore the joystick to its original (centered) position.

We manipulated the task difficulty by controlling three main parameters (independent variables). First, we varied the target’s movement direction by using random angles (uniform distribution) derived from the following angular ranges (AR): ±5°, ±40°, ±75°, ±110°, ±145°, and ±180°. Therefore, the angles (in degrees) were selected randomly for each AR. Second, we used two ‘fixed-direction intervals’ (FDI) of 100 ms and 200 ms to change the target’s movement direction (*i.e.,* the FDI is defined as the interval −consecutive sequence of frames− during which the target’s movement direction remains constant). Therefore, after every FDI, the direction of the target changed with a random angle contained in AR. Finally, we employed multiple target speeds (*v_T_*): 10 °/s, 20 °/s, 30 °/s, 40 °/s, 50 °/s, and 60 °/s (in degrees of visual field per second). The speeds were kept constant within each trial but could also be permuted across trials. In experiments illustrated in *Figures 1,4,* and *5*, we organized different experimental groups of naive participants, assigning each group to solve the tasks at a specific speed (*i.e.,* one speed per group). If, at some moment, the target reached the limits of the arena, we calculated a bounce from the border using the law of reflection as a reference. Thus, the effective reflection angle was equal to the angle at which the moving target was incident on the tangent line to the border of the circular arena, plus the corresponding angular variability given by the AR used for that trial. We discouraged ‘circular evasion tactics’ (turning behavior with relatively constant radius and speed)(Corcoran & Conner, 2016) in escape trials by incorporating two rules into our system: i) we aborted the trial (counting it as an error, and reproducing three brief 500 Hz tones) if the user adopted a circular trajectory involving ≥ 100 frames (*i.e.,* for more than 1.66 s) with either a clock-wise or anti-clock-wise path, and ii) the target’s observed speed (*v_TO_*) was updated every FDI as follows:

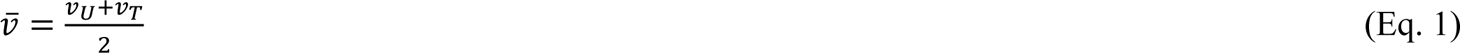

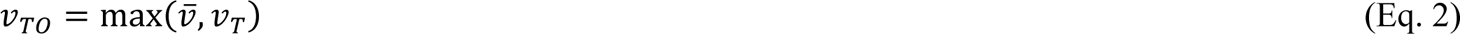

Where *v_U_* is user’s speed, *v_T_* is target’s speed (pre-programmed and fixed during chasing trials), and *v_TO_* is the observed target’s speed after the correction described by Eq. 2. This adjustment, specifically applied to escape trials, allowed *v_TO_* to smoothly align with *v_U_* in situations where the participant moved faster than *v_T_* (as illustrated in *Figure 5c*). Aborted trials (because of turning behavior) were infrequent (≤ 0.01 %/participant) and were not repeated. An entire experimental session lasted less than an hour (Chase trials, FDI = 100 ms, av. trial duration = 2.2 s ± 0.1 s, *n* = 89 participants; FDI = 200 ms, av. trial duration = 2.2 s ± 0.1 s, *n* = 90 participants; Escape trials, FDI = 100 ms, av. trial duration = 3.1 s ± 0.1 s, *n* = 120 participants; Escape trials, FDI = 200 ms, av. trial duration = 3.1 s ± 0.1 s, *n* = 120 participants). In some experiments, we aimed to explore if the observed behavior required visual feedback (Mehta & Schaal, 2002) by using multiple contrasts to represent the participant’s or the target’s dots relative to the 50% gray background of the screen (*Figure 3a*). Other experiments involved masking (*i.e.,* ‘disappearing’) one of the two dots by switching them to 0% relative contrast at different randomly permuted inter-dot distances (IDD, *Figure 3b*). All contrast experiments were preceded by running a routine to linearize the video display monitors using Gamma correct lighting. We also confirmed the reliability of our task with test-retest experiments. There were no restrictions on gaze or head mobility, yet the amount of head movement was minimal (according to our subjective impression). We used programs in MATLAB R2022a (MathWorks, Inc.; Natick, USA) with the Psychophysics Toolbox extensions to run the visuomotor task (PTB-3).

### Analysis

We quantified successful attacks and escapes by detecting whether and when the user and target dots collided with each other externally. We incorporated simulation routines to estimate the probability that these two dots collided by chance (100 iterations/trial). To measure the probability of such random collisions in chase trials, we replayed the observed user’s positions along the trial with a virtual target that moved in a new random trajectory employing the same AR, FDI, and *v_T_* for that trial (*i.e.*, the angular direction changed after each FDI). We used a similar approach to calculate chance levels for successful escapes but based on a complementary AR (labeled AR_C_) when randomizing the target’s motion (*Supplementary Figure 1*). Therefore, we computed a chance level for each trial, lasting the experimentally observed CTs, and scaled this value to match different time windows up to 5 s (as collision risk increases with trial duration). Chance levels depicted in *Figure 1d* were determined based on the durations of the experimental trials. We fitted psychometric curves to successful attack/escape trials using the following equation:

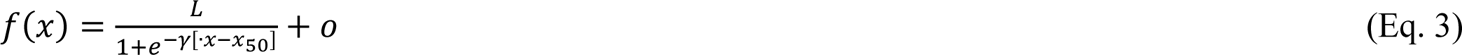

where *L* is the curve’s maximum value, *γ* is the logistic growth rate or slope of the curve, *x_50_* is the *x* value of the sigmoid’s midpoint, and *o* is the offset of the entire curve. Because chasing and escaping trajectories were tracked continuously, we extracted from every frame: the location (in Cartesian and polar coordinates), inter-dot distance (IDD), velocity, and angle between the dots. For chase trials, we also calculated the optimal pursuit angle (given by the imaginary line connecting the participant and target) and an optimal interception angle (*i.e.,* shortest intercept path possible), given by:

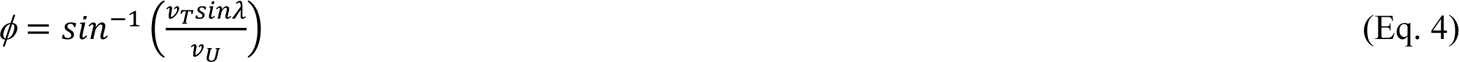

where *v_T_* is the target speed, *v_U_* is the instantaneous speed of the participant, and *λ* is the difference between the tracking angle and the target’s direction (find a detailed description of equations and figures in (Ghose et al., 2006)). If *v_U_* < *v_T_* sin *λ,* then this equation has no solution. However, there are two solutions if *v_U_* > *v_T_* sin *λ*, with only one causing the IDD to decrease (Ghose et al., 2006). To calculate average angles, we first transformed the directions into unit vectors and then calculated the mean resultant vector from which we extracted the mean angular direction. Distributions of pursuit and interception (angular) errors were created by subtracting the participants’ directional angle from the optimal pursuit and interception angles, respectively (all frames from chase trials, *Figure 4b*, and *Supplementary Figure 2*). We calculated %optimal interception angle (%OIA) by using the angular errors obtained on every frame:

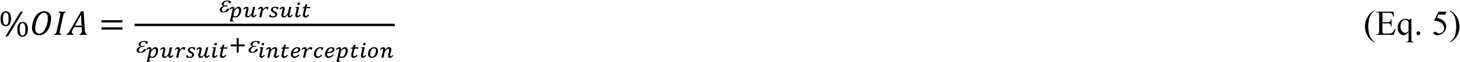

thus, an %OIA greater than 50% implies that user directional angles lean towards interception, indicating a preference for interception behavior. Conversely, a value less than 50% signifies a greater tendency towards tracking behavior. To analyze the angular errors occurring on a circular scale, the pursuit (ε_pursuit_) and interception (ε_interception_) errors from all participant trials were evaluated using the κ (kappa) parameter from the von Mises distribution. This analysis was conducted using CircStat, a MATLAB toolbox designed for circular statistics (Berens, 2009). κ is a measure of concentration for circular data (a reciprocal measure of dispersion; 1/κ is analogous to σ^2^ in the normal distribution). If κ is large, the distribution becomes very concentrated, whereas if κ increases, the distribution approaches a normal distribution with mean μ and variance 1/κ. Lastly, we calculated an averaged %OIA for each participant by taking the reciprocals of the κ values instead of the ‘instantaneous’ ε values, as described in Eq. 5.

### Statistical analysis

We ran an a priori statistical power analysis with the G*Power program (Faul et al., 2009), using an effect size set at 0.2, α at 0.05, and power (1 – β) at 0.9. We found that the minimum sample size to find an effect in our main experiment was ∼60 participants (for experiments illustrated in Figure 1d,e). We used t-tests, non-parametric tests, and Kolmogorov-Smirnov (KS) tests for group and distribution comparisons. Additionally, we employed repeated measures ANOVA (RM-ANOVA) tests with Bonferroni’s or Wilcoxon Signed Rank *post hoc* tests for group comparisons. Two-group comparisons of directional data were performed using the Watson-Williams test, whereas multiple comparisons were made using the multi-sample test for equal median directions, a circular analog to the Kruskal-Wallis test. All statistical analyses were performed using CircStat, a MATLAB toolbox for circular statistics (Berens, 2009), MATLAB R2022 (MathWorks, Inc.; Natick, USA), and STATISTICA (v13.3). We illustrate our group data as averages ± S.E.M. with a significance set at *P* ≤ 0.05. The number of participants used for each task’s analysis is represented inside parentheses in the corresponding figure panels.

## Results

### Design and implementation of a minimalistic visuomotor task

We designed a simple experimental setup to systematically examine visual inputs and motor outputs of the visuomotor system. To achieve this, we developed a computer-controlled visuomotor task to analyze human chase and escape trajectories (*Figure 1a*). In this task, participants used a joystick to move a white dot (third-person viewpoint) that could either chase or escape from a black moving dot controlled by the computer (*i.e.,* the ‘target’; both dots were of 0.3° of visual angle) within a 2D circular arena (*i.e.,* without depth perception). The target traveled at different experimenter-defined velocities (*v_T_*: 10 °/s, 20 °/s, 30 °/s, 40 °/s, 50 °/s, and 60 °/s), with directional changes given by random numbers extracted from uniform distributions covering different angular ranges (AR: ±5°, ±40°, ±75°, ±110°, ±145°, and ±180°). These changes in direction were produced at different ‘fixed-direction intervals’ (FDI: 100, and 200 ms; right panels in *Figure 1a*). Thus, AR controlled the unpredictability of the changes in the direction of the target (computer-controlled black dot), whereas the FDI controlled the frequency of these changes. Before running tests with the task, we conducted a brief pre-training routine to ensure that participants could effectively move the joystick toward static random locations along the border of the arena. Suggesting regular visuomotor abilities, participants quickly improved their performance (Successful hits, from 74.16% ± 1.79% to 99.00% ± 0.41% %; collision time (CT), from 1.65 s ± 0.03 s to 1.15 s ± 0.02 s), reaching their peak in as few as 10 trials (last two trials from pre-training, Kruskal-Wallis test, with Bonferroni *post hoc* test, successful hits, *F* = 1193, *P* > 0.9; CT, *F* = 1181, *P* = 0.11, *n* = 90; *Figure 1b*). The participants’ execution reported in the following sections rendered stable performance metrics during the entire experimental sessions without signs of being affected by fatigue or other factors (sample plots in *Figure 1c*).

### Uncertainty in path direction and the velocity of the moving target determine task performance

While the task we created allows for the exploration of various visuomotor strategies, this investigation aimed to provide a proof-of-concept and demonstrate the platform’s capability to capture and analyze visuomotor behaviors under controlled uncertainty. Because our visuomotor task involves a target moving in trajectories with variable predictability, we first explored how key task parameters influenced the participants’ performance (Domenici et al., 2011). We reasoned that solving the task should involve integrating relational information such as position and velocity. For each individual, we measured the number of successful attacks and escape trials, and their CTs (max. trial duration of 5 s, see **Method**). We created twelve permuted groups of participants tested with specific target velocities (*v_T_*: 10 °/s, 20 °/s, 30 °/s, 40 °/s, 50 °/s, and 60 °/s), and FDI (100 ms, and 200 ms) values (*i.e.,* 6 groups of 15 participants per *v_T_*\*FDI, see **Method**). We illustrate the group averaged data from these experiments in *Figure 1d,e*. AR exerted control over the variability in the target’s trajectory, introducing uncertainty in its position and movement. Therefore, increasing AR reduced successful attacks in chasing trials (Kruskal-Wallis test, ±5° *vs.* ±180°, FDI = 100 ms, successful attack, *F* = 5.58, *P* = 0.018; CT, *F* = 10.31, *P* < 0.002; FDI = 200 ms, successful attack, *F* = 5.37, *P* = 0.02; CT, *F* = 8.13, *P* = 0.004, *Figure 1d*) but improved successful escapes (FDI = 100 ms, successful escape, *F* = 78.55, *P* < 0.001; CT, *F* = 37.40, *P* < 0.001; FDI = 200 ms, successful escape, *F* = 67.10, P < 0.001; CT, *F* = 37. 90, P < 0.001, *Figure 1e*). In contrast, slower *v_T_* values improved successful attacks (Kruskal-Wallis Multicomparison test, FDI = 100 ms, successful attack, *F* = 56.07, *P* < 0.001; CT, *F* = 30.84, *P* < 0.001; FDI = 200 ms, successful attack, *F* = 65.88, *P* < 0.001; CT, *F* = 48.19, *P* < 0.001, *Figure 1d*), and escapes (FDI = 100 ms, successful escape, *F* = 62.47, *P* < 0.001; CT, *F* = 79.31, *P* < 0.001; FDI = 200 ms, successful escape, *F* = 51.10, *P* < 0.001; CT, *F* = 61.79, *P* < 0.001, *Figure 1e*).

The dependency of task performance on AR and *v_T_* was well described by psychometric curves (continuous colored lines in *Figure 1d,e*). Both the offsets (Chasing trials, FDI = 100 ms, m = −0.01, *P* < 0.001; FDI = 200 ms, m = −0.01, *P* < 0.001; Escaping trials, FDI = 100 ms, m = −0.01, *P* < 0.001; FDI = 200 ms, m = −0.007, *P* < 0.01), and the area under these psychometric curves (AUC; Chasing trials, FDI = 100 ms, m = −0.06, *P* < 0.001; FDI = 200 ms, m = −0.06, *P* < 0.001; Escaping trials, FDI = 100 ms, m = −0.06, *P* < 0.001; FDI = 200 ms, m = −0.03, *P* < 0.03) decreased with *v_T_*, confirming that higher *v_T_* values led to reduced averaged performance (insets in *Figure 1d,e*). Parameter fits from the psychometric curves and their regressions against task parameters are shown in *Supplementary Tables 1-3*. We conducted a multi-factor analysis of variance and confirmed that AR (Chasing trials, *F* = 16.78, *P* < 0.001; Escaping trials, *F* = 70.87, *P* < 0.001), FDI (Chasing trials, *F* = 9.69, *P* = 0.001; Escaping trials, *F* = 4.37, *P* = 0.03), and *v_T_* (Chasing trials, *F* = 685.51, *P* < 0.001; Escaping trials, *F* = 225.51, *P* < 0.001) influenced successful attacks and escapes, with relevant interactions between AR and *v_T_* (Chasing trials, *F* = 1.69, *P* < 0.02; Escaping trials, *F* = 1.70, *P* = 0.02), and between FDI and *v_T_* (Escaping trials, *F* = 28.64, *P* < 0.001). There were similar dependencies for the observed CTs (Chasing trials, AR : *F* = 7.41, *P* < 0.001; FDI : *F* = 6.76, *P* < 0.001; *v_T_* : *F* = 80.18, *P* < 0.001; AR\**v_T_* : *F* = 2.13, *P* = 0.001; Escaping trials, AR : *F* = 146.98, *P* < 0.001; FDI : *F* = 0.77, *P* = 0.38; *v_T_* : *F* = 312.56, *P* < 0.001; AR\**v_T_* : *F* = 6.13, *P* < 0.001; FDI\**v_T_* : *F* = 25.43, *P* < 0.001). *v_T_* had the biggest effect size for chasing and escaping trials (Chasing trials, AR = 0.11, TDI = 0.19, *v_T_* = 0.96; Escaping trials, AR = 0.15, FDI = 0.14, *v_T_* = 0.39). In *Supplementary Figure 1*, we describe and illustrate how we measured the probability of random collisions to estimate the chance levels depicted in *Figure 1d*.

Because FDI controls the frequency of changes in the target’s direction, we reasoned that it could indirectly impact task uncertainty and performance. Intuitively, we reasoned that higher FDI values could lead to higher successful attacks but reduced escapes. Using 100% contrast for both user and target, and a constant target speed of *v_T_* = 40 °/s, we tested the effects on task performance of randomly permuting FDI values (FDI: 100 ms, 125 ms, 150 ms, 175 ms, and 200 ms). The FDI values tested had no apparent influence on successful attacks and escapes (AR = ± 5°, *P* > 0.05; AR = ± 40°, *P* > 0.50; AR = 110°, *P* > 0.50; *n* = 20 per group, *Figure 1f*), with similar results on CT, and *v_U_* (not illustrated). In other words, participants showed a remarkably stable performance despite the multiple FDI values tested.

### Task performance is consistent and reliable on a wide range of experimental conditions

Next, we explored how the participants adjusted their performance to more challenging and volatile conditions. More specifically, we tested whether chasing/escaping performances were sensitive to sequential changes in parameter volatility during the experimental session (Trevino, Castiello, et al., 2021). Stochastic feedback control theory suggests that goal-directed corrections should occur when external noise interferes with task performance (Todorov & Jordan, 2002). By using an FDI = 100 ms, *v_T_* = 40 °/s, and 100% contrast, we employed five blocks of graded AR variance by increasing/decreasing the bounds of the uniform distributions (±0°, ±22.5°, ±45°, ±67.5°, ±90°) around an average change in the direction of the moving target of ±45°. We trained two groups of naïve participants, with increasing (gray) and decreasing (pink) AR variance. Notably, we found no differences in their chasing performance both during training (5 blocks of 50 trials, Successful attacks: *F* < 35, *P* > 0.55; CT: *F* < 49, *P* > 0.48), and when testing both groups with identical AR conditions at the end of training (6^th^ block, last 50 trials: Successful attacks: *F* = 0.39, *P* = 0.55; CT, *F* = 2.45, *P* = 0.11; upper panels in *Figure 2a*). Similarly, there were no group differences in escape trials (5 blocks of 50 trials, successful escapes: *F* < 2.68, *P* > 0.21; CT: *F* < 2.56, *P* > 0.15; 6^th^ block: Successful attacks: *F* = 0.01, *P* = 0.91; CT, *F* = 0.27, *P* = 0.60). We also explored increasing/decreasing the FDI variance in two new groups of participants but found similar performance between groups during training (5 blocks of 50 trials, Successful attacks: *F* < 1.56, *P* > 0.21; CT: *F* < 1.85, *P* > 0.17; successful escapes: *F* < 11.95, *P* > 0.11; CT: *F* < 2.16, *P* > 0.14), and when testing them with identical FDI conditions at the end of training (6^th^ block, last 50 trials: successful attacks: *F* = 2.77, *P* = 0.09; CT: *F* = 3.57, *P* = 0.06; successful escapes: *F* = 0.11, *P* = 0.73; CT: *F* = 0.82, *P* = 0.36; lower panels in *Figure 2b*). These results suggest the involvement of task-specific control laws that prioritize correcting motor errors to preserve performance (Ernst & Banks, 2002; Todorov & Jordan, 2002). Hence the observed ‘immunity’ of performance to the volatility introduced in the target’s kinematics.

**Figure 2.**
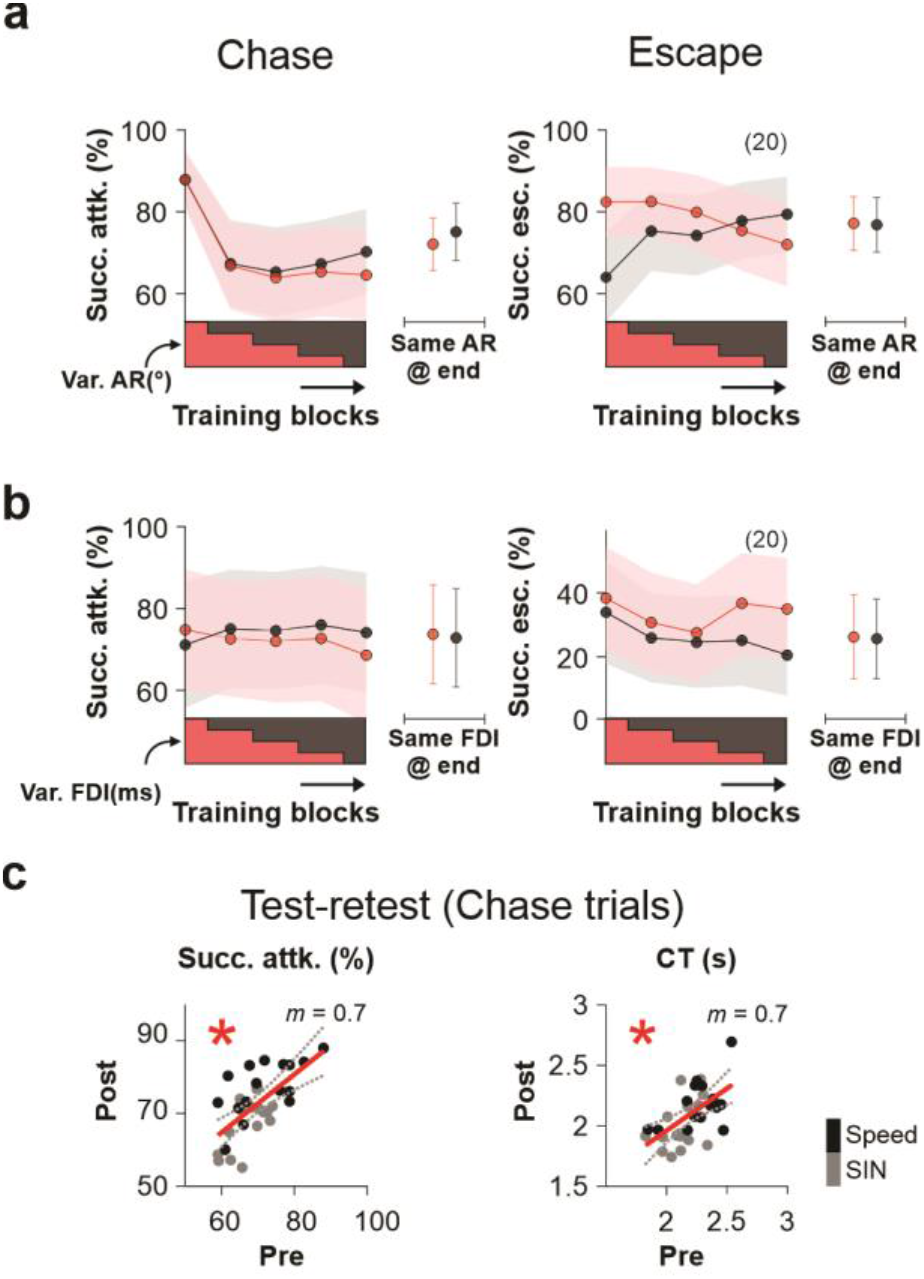
The visuomotor task renders a stable chasing/escaping performance across various experimental conditions. **a** Chasing (left) and escaping (right) performance from two groups of participants trained with increasing (gray) and decreasing (pink) AR variance (blocks of 100 trials, bounds from 0 to ±90°). The sixth block, represents av. performance (tested with identical AR conditions for both groups) after completing training with the ascending/descending ramps. **b** Similar experiment as in previous panel, but changing the FDI variance. **c** Scatter dots test-retest experiments with linear regressions for chasing performance and corresponding av. CTs. Colors of the dots represent different experimental procedures: one in which different *v_T_* values were randomly permuted (’Speed’ experiments), and one in which they were changed using a sinusoidal function (’SIN’ experiments). Number of participants per experiment in parentheses. Asterisks depict significant slopes (*P* ≤ 0.05).

Motivated by the behavioral stability observed in the previous results, we assessed the reliability of the visuomotor task by comparing the performance of a new group of participants across two different dates (test re-test procedure; days between tests: 24.81 days ± 0.97 days, mode: 26, min: 13, max: 37, *n* = 15). We found similar performance during both tests (successful attacks, Pre: 70.04 % ± 0.01 %, post: 72.73 % ± 0.02 %, paired t-test, *P* = 0.06; CT, Pre: 2.17 s ± 0.03 s, post: 2.13 s ± 0.04 s, paired t-test, *P*= 0.14), and a Pearson correlation coefficient with acceptable reliability (Hits, correlation coefficient: 0.773, *P* < 0.001; *Figure 2c*). Such stable behavioral differences across participants suggest that cognitive and executive processes could underlie such differences.

Because the visual system is susceptible to changes in spatial information, we reasoned that stimulus detectability should affect movement precision and optimal interception (but see Schroeger et al., 2021). First, we measured the impact of varying the contrast of either the participant (white dot), or the target (black dot) relative to the 50% gray background arena. Notably, reducing the participants’ contrast did not affect their chasing performance (Successful attack, m = 6.71 × 10^-6^, *P* = 0.99; CT, m = 3.55 × 10^-4^, *P* = 0.77; blue dots in left panel from *Figure 3a*). However, reducing the contrast of the moving target impaired attacking performance (Successful attack, m = 0.003, *P* < 0.001; CT, m = 9.03 × 10^-4^, *P* = 0.59, black dots in left panel from *Figure 3a*). The opposite happened in escape trials (changes in user contrast, blue dots, Successful escapes, m = −7.03 × 10^-4^, *P* = 0.02; CT: m = 0.0025, *P* = 0.02; changes in target’s contrast, black dots, Successful escapes, m = 9.73 × 10^-5^, *P* = 0.73; CT, m = 0.002, *P* = 0.06; right panel from *Figure 3a*). These results indicate that participants used proprioceptive information to solve the task but required visualization of the moving target to chase and escape (Franklin & Wolpert, 2011). Also, the lack of visual information about the user did not impair the control of hand-movement velocity (Hocherman & Levy, 2000).

**Figure 3.**
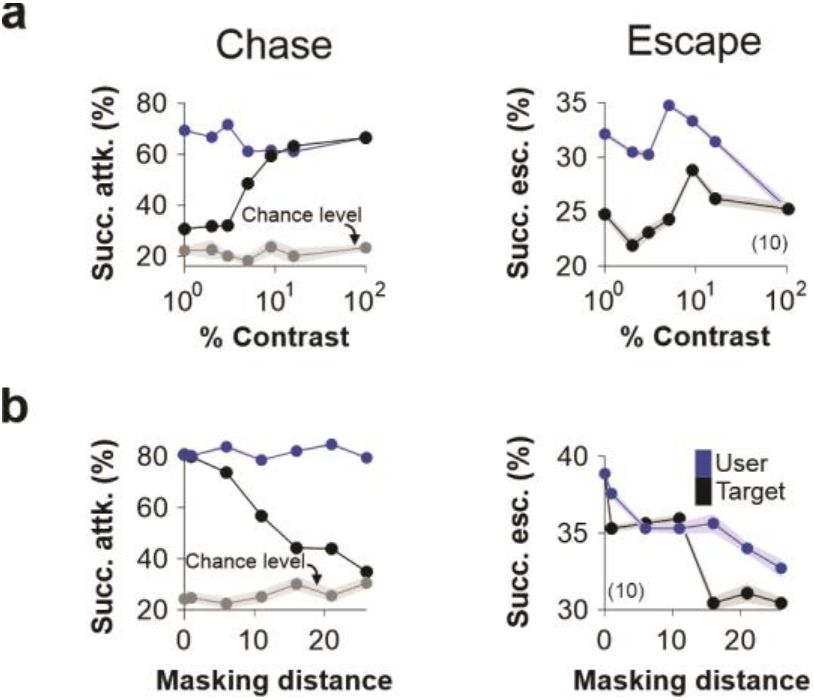
The visuomotor task involves visual and propioceptive signals. **a** Succesful attacks (left) and escapes (right) as a function of changes in user (blue dots) or target’s (black dots) contrast (FDI = 100 ms; AR = ± 75°). **b** Attack (left) and escape (right) performance as a function of the inter-dot distance (IDD) that we used to blank-out (mask) the user, or the target (same color coding as previous panel). Chance level in gray. All our figures show group data as averages ± S.E.M. Number of participants per experiment in parentheses.

We next explored whether and how masking (*i.e.,* stimuli switched to a 0% relative contrast against the background) the participant’s (white dot) or the target’s dot (black dot) at different inter-dot distances (IDD, the Euclidean distance between participant and moving target (Mehta & Schaal, 2002)) changed task performance. In agreement with our previous findings, chasing performance dropped with increasing masking distances (masked target −in black−, Successful attacks, m = −0.01, *P* < 0.001; CT, m = −0.006, *P* = 0.43; masked participant −in blue−, Successful attacks, m = −0.0001, *P* = 0.81; CT, m = −0.014, *P* = 0.11), but the opposite happened in escaping trials (masked target −in black−, Successful escapes, m = −0.002, *P* < 0.008; CT, m = −0.003, *P* = 0.23; Masked participant −in blue−, Successful escapes, m = −0.0002, *P* < 0.003; CT, m = −0.0002, *P* = 0.93; *Figure 3b*). These results reveal that the visuomotor task can be solved with graded efficiencies using predictive signals and/or predicted states despite the reduced information about the moving target.

### Identifying interception behavior during chasing trials

Chasing a moving target can combine pursuit and interception actions (Ghose et al., 2006; Tsutsui et al., 2020; Zhao & Warren, 2015). We built a geometrical model of potential chaser/escaper interactions to quantify the relative contribution of these two strategies when participants chased the moving target (*Figure 4a*; see **Method**). The attacker’s direction, *v_U_*, *v_T_*, and the target’s movement direction were continuously measured. With this information, we calculated the optimal pursuit (sky blue) and interception (hot pink) angles for every acquired frame (*Figure 4a*). We then calculated the (angular) pursuit and interception errors, respectively (*ε_pursuit_* = the angular difference between the current chaser’s direction and the optimal pursuit angle; *ε_interception_ =* the angular difference between the current chaser’s direction and the optimal interception angle). We extracted the concentration parameter (κ, kappa, a reciprocal measure of dispersion: 1/κ is analogous to σ^2^ in the normal distribution, see **Method**), and measured the area under the curve (AUC, using the raw group probabilities) from these angular error distributions. Because interception becomes more challenging with faster targets, we reasoned that the angular errors should increase with *v_T_* (*i.e.,* broader error distributions, smaller κ). Sample frequency distributions in *Figure 4b* (from experiments with FDI = 100 ms, AR = ± 40°) illustrate how these error distributions dispersed with *v_T_* (the complete set of panels is shown in *Supplementary Figure 2*). Notably, AR dispersed the *ε_pursuit_* distributions but had virtually no effect on the shape of the *ε_interception_* distributions (*v_T_* = 30 °/s; *Figure 4c*). Therefore, *v_T_* was the main factor influencing the dispersion of the angular error distributions (FDI = 100 ms; κ_pursuit_, m = 0.44, *P* = 0.01; AUC, m = 2.21 × 10^-5^, *P* = 0.03; κ_interception_, m = 0.28, *P* = 0.07; AUC, m = −3.91 × 10^-5^, *P* = 0.007; *Figure 4d,e*).

**Figure 4.**
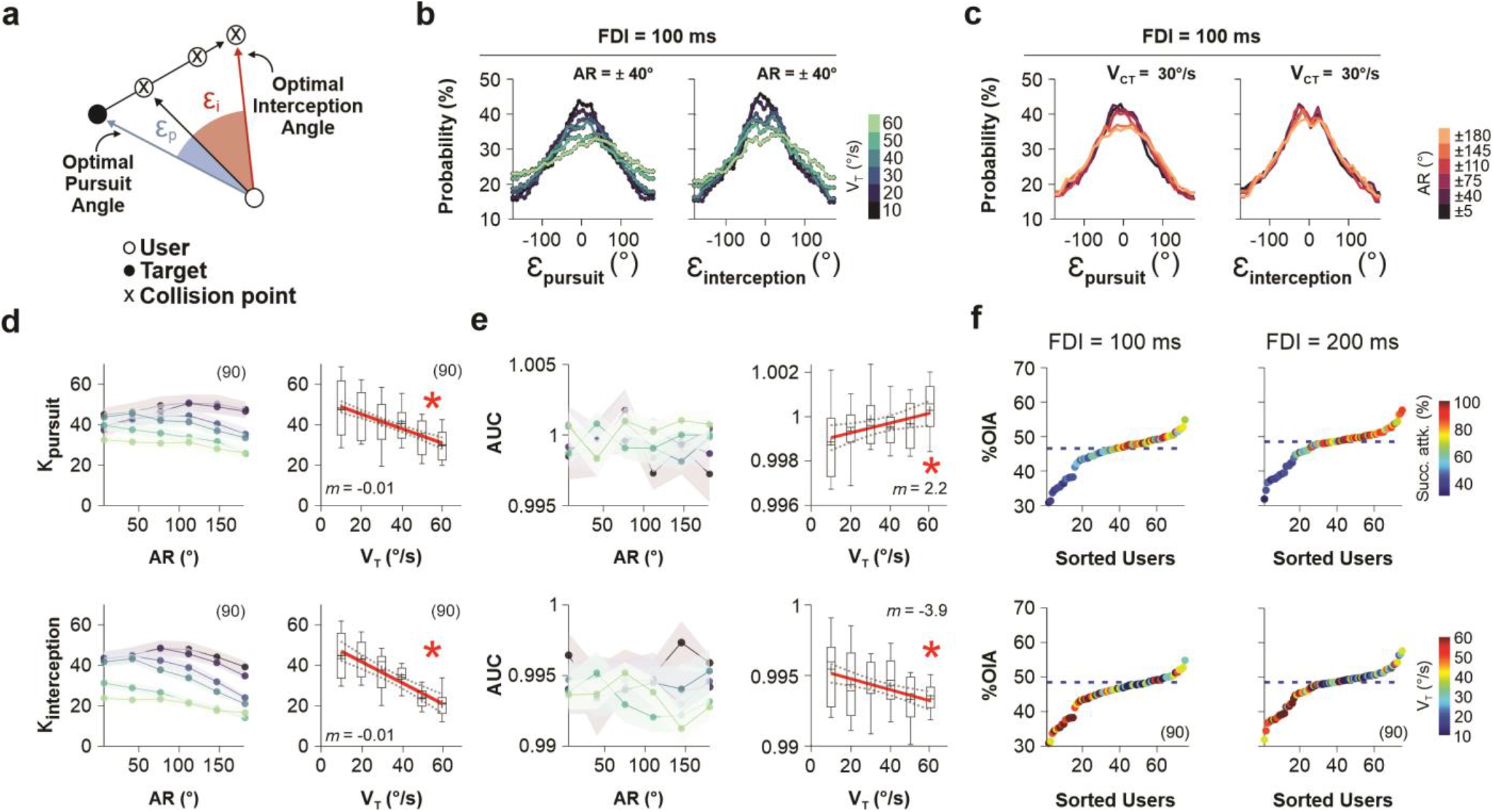
Assessing tracking and interception strategies during chasing trials. **a** Geometrical model to characterize how much the participants were adopting either a pursuit or an intercepting strategy. For each frame captured (approximately every ∼16.66 ms), two angular errors were calculated: the angular pursuit error (ε_pursuit_) represented by the optimal pursuit angle minus the participant’s current direction (shown in sky blue), and the angular interception error (ε_interception_) represented by the optimal interception angle minus the current direction (shown in hot pink; see **Method**). By analyzing these angular errors, we can infer the dominant strategy employed by the participant. If a participant predominantly adopts one strategy over the other, the corresponding angular errors for that strategy are expected to be smaller, while the complementary strategy would exhibit larger errors. **b** Sample group frequency distributions of pursuit (left) and interception (right) angular errors (complete analysis is shown in *Supplementary Figure 2*). Target computer speeds (*v_T_* depicted in the colorbar), determine the shape of the distributions. **c** AR affects the amplitude of the ε_pursuit_ but not the ε_interception_ distributions. **d** Average concentration parameter, κ (kappa, from the von Mises distribution, see **Method**), calculated from the angular errors from each participant as a function of AR. Whisker plots to the right show κ as a function of *v_T_*. **e** Area under the curve (AUC) from these frequency distributions as a function of AR. Same color-coding for *v_T_* as in panel **b**. Whisker plots to the right show AUC as a function of *v_T_*. **f** Average %OIA adopted by each participant by comparing ε_pursuit_ and ε_interception_ (see **Method**). Dots represent participants, sorted from lowest to highest interceptors. Color of the dots represents chasing performance (upper panel), or *v_T_* (lower panel). Number of participants per experiment in parentheses.

By extracting the κ_pursuit_ and κ_interception_, we assessed the average %optimal interception angle (%OIA) adopted by each participant (see **Method**). An average %OIA < 50% indicates that the participant tended towards using a pursuit strategy, pointing the joystick towards the center of the moving target, whereas an %OIA > 50% indicates that the participant employed more of an interception trajectory, pointing towards the expected future position of the moving target. The sorted averaged %OIA (from all the trials from each participant) is shown in *Figure 4f*. By employing a median split, participants were categorized into ‘low’ or ‘high’ interceptors, revealing that ‘high’ interceptors exhibited higher average performance (upper panels) but smaller average *v_T_* (lower panels; FDI = 100 ms, Successful attacks below median 48.93 % ± 1.56 %; av. *v_T_* below median 49.47 °/s ± 1.66 °/s; Successful attacks above median 79.22 % ± 2.37 %; av. *v_T_* above median 26.21 °/s ± 2.49 °/s; *P* < 0.001; FDI = 200 ms, Successful attacks below median 55.12 % ± 2.75 %; av. *v_T_* below median 44.44 °/s ± 2.43 °/s; Successful attacks above median 83.37 % ± 1.82 %; av. *v_T_* above median 25.55 °/s ± 1.86 °/s; *P* < 0.001). ‘High’ interceptors also exhibited higher performance variance compared to ‘low’ interceptors (FDI = 100 ms, Variance in successful attacks below median 0.0090 Variance in successful attacks above median 0.0208; Variance in *v_T_* below median 102.41; Variance in *v_T_* above median 229.72; FDI = 200 ms, Variance in successful attacks below median 0.0333; Variance in successful attacks above median 0.0145; Variance in *v_T_* below median 261.61; Variance in *v_T_* above median 152.52). These results suggest that ‘high’ interceptors may prioritize accuracy in their interceptive actions, whereas ‘low’ interceptors demonstrate more consistent performance, indicating a narrower range of variability in their interceptive abilities. These results also suggest that individuals with different interception profiles may employ distinct strategies (or have varying proficiency levels) in intercepting moving targets. The lower plots in *Figure 4f* indicate that %OIA values above the median tend to occur with a *v_T_* of approximately 30 °/s.

### Fast adjustments in interception trajectories within trials

Pursuit and interception strategies could coexist in our trials depending on the time window participants had to elaborate their chasing plan (Dessing et al., 2005). We hypothesized that participants adjust their interception strategy as they approximate the target. Therefore, to search for dynamic changes in %OIA, we analyzed the last ∼800 ms of the chasing trials before the user collided with the moving target (*i.e.,* event-triggered averages). We calculated the group frequency distributions of observed *ε_pursuit_* and *ε_interception_* (0 represents the optimal pursuit/interception angle) and how these angular error distributions evolved (*i.e.,* got narrower) as the participants got closer to the target (FDI = 100 and 200 ms pooled together, *Figure 5a*). This analysis reveals that the participants increased their interception strategy (%OIA) as they got closer to the target (smaller IDDs), just before the collision. These results illustrate how our method is robust in detecting an increase in the adoption of the interception strategy compared to the pursuit strategy during the final phase of chasing trials, prior to collision.

**Figure 5.**
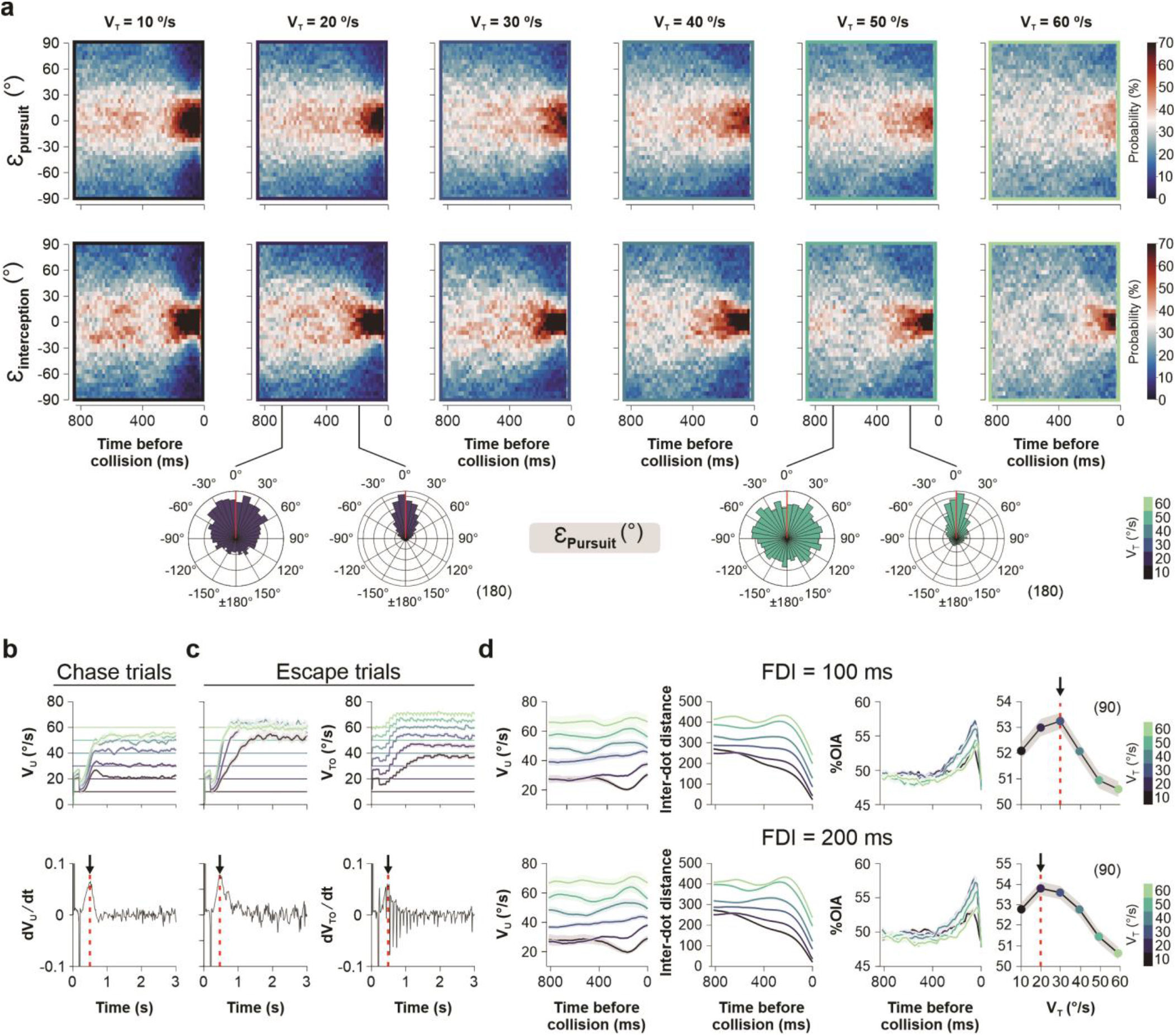
Dynamic changes in pursuit and interception strategies during chasing trials. **a** Probability distributions of the pursuit and interception angular errors as a function of the number of frames before colliding with the target (*i.e.,* time before collision). Panels are arranged from left to right using lowest to highest *v_T_*. Polar panels below the colormaps show the distributions of angular interception errors at two intervals before collision. The probability distributions were obtained from pooling together data from experiments performed with FDI = 100 and 200 ms. **b,c** Speed analysis for chasing and escaping trials, respectively. The upper panels show average *v_U_* and *v_C_* over time within trials. *v_T_* values are depicted as horizontal thin lines. Panels below show the group average time derivative of the upper traces. **d** Collision triggered averages of user speed (*v_U_*), inter-dot distance (IDD), and %Interception strategy before colliding with the target (upper panels: FDI = 100 ms; lower panels: FDI = 200 ms). Fourth column of panels show the %Interception strategy (*i.e.,* %OIA) as a function of *v_T_*, depicted in the colorbar. Number of participants per experiment in parentheses.

Multiple factors affect moving targets’ interception (Brenner & Smeets, 2015). In our case, the amount of %OIA employed could depend on experimentally controlled factors like *v_T_*. Indeed, due to sensory and motor delays, target interception requires a continuous forecast of the target’s future location based on its current position and speed (Soechting et al., 2009). Furthermore, the impact of temporal errors can be reduced by approaching the target with a velocity that matches its speed (*i.e., v_U_ → v_T_*). Thus, we explored whether the participants adjusted their speed (*v_U_*) to *v_T_*, improving the odds of a successful attack/escape (Corcoran & Conner, 2016; Domenici et al., 2011; Howland, 1974). We conducted a new set of experiments with randomly permuted *v_T_* values across trials (10°/s, 20°/s, 30°/s, 40°/s, 50°/s, 60°/s) and found that participants quickly adjusted their *v_U_* to match *v_T_*, both in chase (*Figure 5b*) and escape (*Figure 5c*) trials. Notably, these *v_U_* adjustments occurred relatively fast within the first ∼650 ms of the trial, as revealed by analyzing the average of the time derivatives of *v_U_* from all *v_T_* tested (lower panels in *Figure 5b,c*). This implies that the quick *v_U_* adjustments at the beginning of the trials cannot directly explain the changes in %OIA observed during the last ∼400 ms before collision (chase trials, FDI = 100 ms, av. trial duration = 2.25 s ± 0.02 s, *n* = 90 participants; Chase trials, FDI = 200 ms, av. trial duration = 2.20 s ± 0.02 s, *n* = 90 participants). To confirm this idea, we explored the evolution of IDD, *v_U_*, and %OIA during the last 800 ms before participants effectively collided with the moving target (*Figure 5d*). The averaged traces showed stable *v_U_* values during all these frames before the collision. However, the IDD dropped faster with slower *v_T_* values (panels on the second column), and the %OIA increased (up to ∼9%) as the user got closer to the target (panels on the third column), with peak %OIA with a *v_T_* of 20-30°/s (panels on the fourth column of *Figure 5d*). These results indicate that *v_T_*, but not the quick adjustments in *v_U_* within trials, influenced the late interception strategy employed. Based on these results, selecting a vT of 30°/s for future experiments is recommended, as it would enrich the trials with the interception strategy.

### Developmental regulation of chasing and escaping performance

Because our data showed distinctive but stable behavioral metrics across participants (*Figure 2c*), one interesting possibility is that chasing/escaping measures could also reflect traits that develop and improve with age (Trevino, Beltran-Navarro, et al., 2021). We thus tested our task in three groups of children/youngsters with ages of 6 (6.65 ± 0.23 years old, *n* = 20), 10 (10.35 ± 0.23 years old, *n* = 20), and 20 (21.72 ± 0.33 years old, *n* = 20) years, respectively (cross-sectional approach). We collected their brief clinical histories and employed guidelines of typical development as part of our inclusion criteria (see **Method**). We used a shorter version of the task (*Figure 6a*), only involving 120 trials per participant, with a *v_T_* = 20 °/s and AR = ± 5° and ± 180°. There were robust group differences in chasing and escaping performance across all ages (Successful attacks, AR = ± 5°, *F* = 32.14, *P* < 0.001; AR = ± 180°, *F* = 32.14, *P* < 0.001; Successful escapes, AR = 5°, *F* = 28.36, *P* < 0.001; AR = ± 180°, *F* = 20.90, *P* < 0.001; upper panels in *Figure 6b,c*), and all these responses differed from chance levels (*P* < 0.012 for all cases; gray whisker plots). CTs also exhibited a clear reduction through development (Successful attacks, AR = ± 5°, *F* = 33.03, *P* < 0.001; AR = ± 180°, *F* = 33.25, *P* < 0.001; Successful escapes, AR = ± 5°, *F* = 38.71, *P* < 0.001; AR = ± 180°, *F* = 31.32, *P* < 0.001; lower panels in *Figure 6b,c*). Therefore, as children grew, their chasing/escaping performance improved, demonstrating that our task is sensitive to age-related performance changes, making it suitable for future developmental studies.

**Figure 6.**
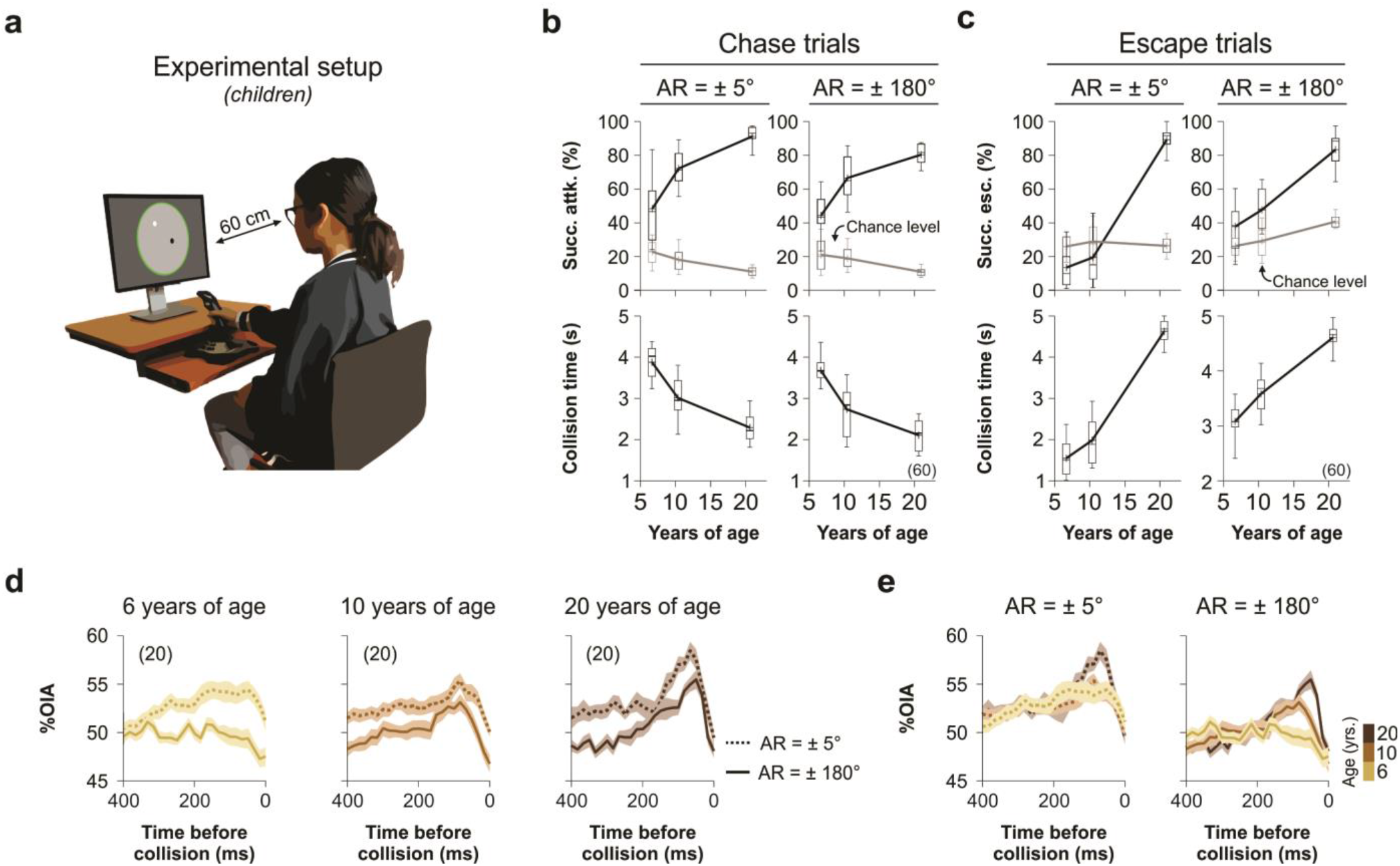
Changes in chasing and escaping performance through development. **a** Cartoon of the experimental setup for children. **b,c** Whisker plots of successful attacks (left) and escapes (right) from children and youngsters in three groups: 6, 10 and 20 years of age. Panels are separated for AR = ±5°, and AR = ±180°, with the inferior plots showing the collision times (CTs). Gray whisker plots correspond to chance levels. **d** Collision triggered averages of %Interception strategy as a function of the last fifty frames before collision (*i.e.,* time before collision; complete figure in *Supplementary Figure 3*). Panels are arranged as a function of age group. **e** Same analysis but with overimposed traces for the different age groups, which are represented in the colorbar to the right. Number of participants in parentheses.

Next, we estimated the %OIA used by the children during the last 400 ms of their chasing trials before effectively colliding with the target (same procedure as in the previous section). Using an ANOVA multicomparison test involving the two AR values (± 5°, and ± 180°) and three age groups (6, 10, and 20 years of age), we found that *v_U_* (last 20 frames ≈ 333 ms, fixed *v_T_* = 20 °/s, *F* = 104.69, *P* < 0.001), chasing IDD (*F* = 592.16, *P* < 0.001), and %OIA (*F* = 7.7, *P* < 0.001) were influenced by age (better performance with age; *Figure 6d* and *Supplementary Figure 3a*). Similarly, *v_U_* (, *F* = 89.39, *P* < 0.001), chasing IDD (*F* = 28.10, *P* < 0.001), and %OIA (*F* = 422, *P* < 0.001) were all sensitive to AR (*Figure 6e* and *Supplementary Figure 3b*). Chasing IDD showed the strongest interaction between AR and age (*F* = 14.97, *P* < 0.001). These findings suggest an age-related increase in the late interception strategy.

### Chasing efficiency is determined by the amount of interception strategy employed

Chasing trajectories that get closer to the moving target increase the odds of producing successful collisions. Because the area under the IDD trace (prior to collision) serves as a proxy for such chasing efficiency (η, *i.e.,* efficient chasers have smaller IDD areas), we wondered whether η could predict the amount of %OIA employed before the collision. Taking the data from the 90 adult participants that participated in our main experiment (AR = ± 5°, ± 40°, ± 75°, ± 110°, ± 145°, ± 180°; FDI = 100 ms, 200 ms; *v_T_* = 10°/s, 20°/s, 30°/s, 40°/s, 50°/s, 60°/s), we created two subgroups by using a median split of the area under the averaged IDD traces (AUC; from each participant, ranging from 416 ms to 50 ms before colliding with the target). Once tagging ‘inferior’ (deep yellow) and ‘superior’ chasers (dark green), we calculated the corresponding averaged %OIA traces before the collision. The right panel in *Figure 7a* confirms that the chasing efficiency can be directly linked to participants using more content of %OIA (*F* = 5.34, *P* = 0.02). We performed a similar analysis with the children’s data by splitting the observed distributions into three equal parts: ‘lower’ (deep yellow), ‘medium’ (pink), and ‘higher’ (dark green) performers (*Figure 7b*). Once again, higher performers showed a higher content of a late interception strategy (*F* ≥ 19, *P* ≤ 0.001, for all cases).

**Figure 7.**
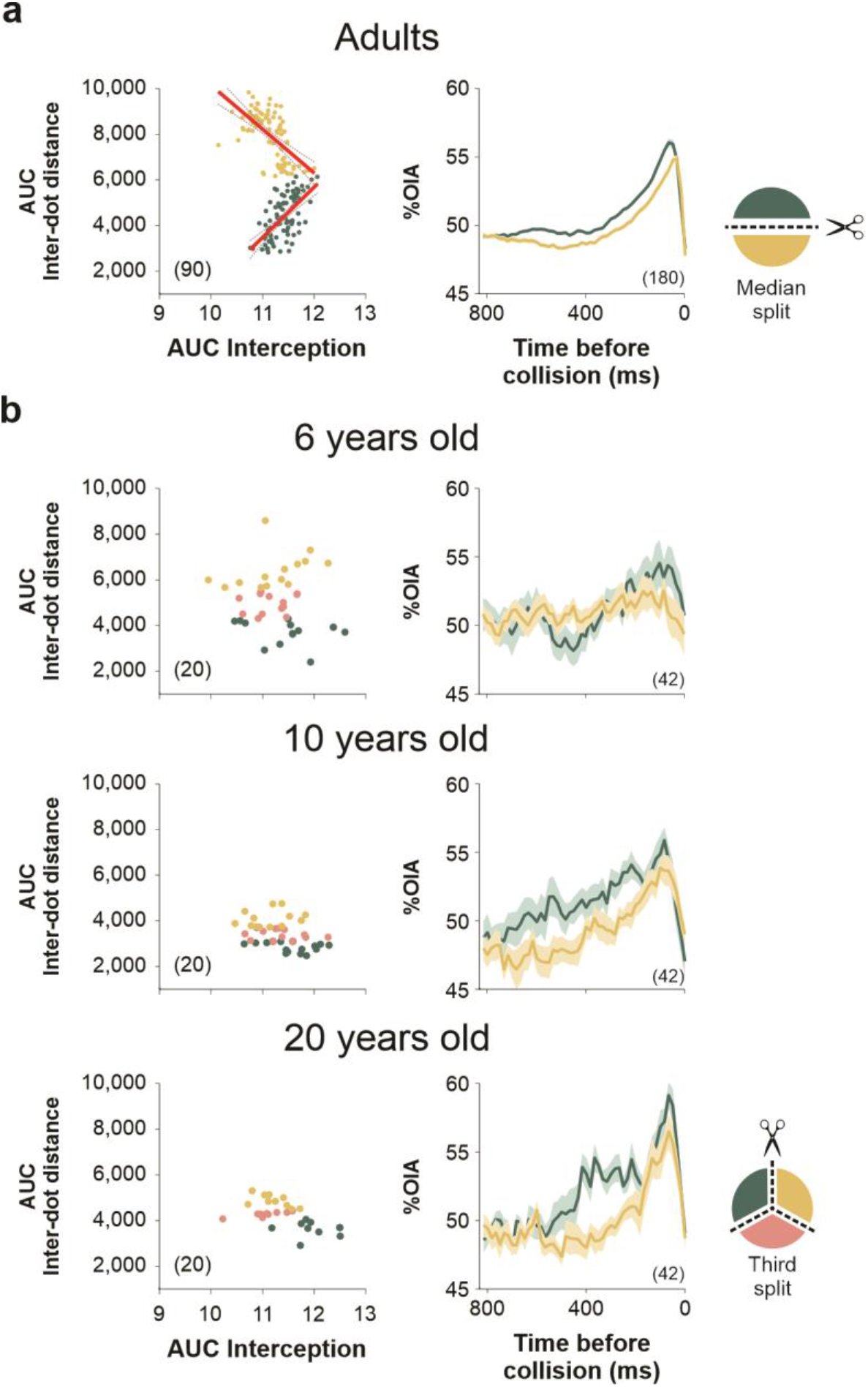
Chasing efficiency depends on the amount of interception strategy employed. **a** Left panel shows the median split into ‘inferior’ (deep yellow) and ‘superior’ (dark green) adult chasers by calculating the area under the curve (AUC) of IDD (from 416 ms to 50 ms before collision). The right panel shows the average %OIA for ‘inferior’, and ‘superior’ chasers. **b** Similar analysis applied to children data by sorting them into ‘inferior’ (deep yellow), ‘median’ (pink), and ‘superior’ (dark green) chasers. Panels on the right show the averaged %OIA traces for ‘inferior’ and ‘superior’ chasers only. Number of participants per age group in parentheses.

## General Discussion

We created a minimalistic computer task that allowed participants to interact with a moving dot on a computer screen. A novel feature of our task is that the target moved in random and unprescribed rectilinear trajectories with different amounts of path uncertainty (AR) and speeds (*v_T_*). To solve the task, users had to either chase or escape from the moving dot using a joystick (*i.e.,* manual control). They became acquainted with the mechanical impedance of the joystick (*i.e.,* resistance to motion) during a brief pre-training phase (Franklin & Wolpert, 2011; Kording & Wolpert, 2004), which also served to confirm that they had proper visuomotor abilities (Mehta & Schaal, 2002). Their performance was stable and did not improve (or worsen) during the experimental sessions. The task gave us parametric control of the unpredictability of the target’s movement direction, how frequently we changed it’s direction, and it’s speed (distance and location estimates worsen with speed). Each task parameter (independent variables) had an associated controlled variability, leading to uncertainty in the predicted target position. Therefore, participants had to use a continuous sensory feedback to adapt to the statistical fluctuations we imposed for each trial (Clark, 2016; Kording & Wolpert, 2004; Piray & Daw, 2020). The responses we characterized in our task could not be observed by chance, and AR and *v_T_* influenced successful attacks and escapes (Coudiere et al., 2022; Domenici et al., 2011; Hocherman & Levy, 200^0^; Lam & Zenon, 2021). Higher AR and *v_T_* values in chasing trials increased uncertainty in the escaping path, making it more challenging for users to reach the target. Conversely, increasing AR but decreasing *v_T_* during escape trials reduced the efficiency of target tracking, thereby enhancing the participants’ probability of successful escape. Assuming that participants used the same control strategy to solve all the trials (Todorov & Jordan, 2002), this would imply that AR and *v_T_* directly influenced task difficulty. Chasing and escaping performances were well described with psychometric curves.

Both adaptability and stability complement human (and animal) behavior (Behrens et al., 2007; Piray & Daw, 2020; Warren, 2006). Nevertheless, action dynamics are non-stationary, changing as the agent interacts with the environment (Warren, 2006). Studies show that sensory uncertainty can increase biases in perceptual judgments (Kwon & Knill, 2013; Trevino et al., 2020). We tested blocks with monotonic changes in AR and FDI variance. Intuitively, higher input volatility should lead to higher outcome variance (Piray & Daw, 2020). However, we found no difference between groups trained with positive and negative variance gradients, indicating that participants’ performance was stable and resistant to our manipulations in parameter variance. These results align with the view that the motor system optimizes performance by incorporating sensory and motor uncertainty (Todorov & Jordan, 2002). It adapts motor behavior to varying levels of uncertainty, demonstrating efficient Bayesian processing (Knill & Pouget, 2004). Similarly, predictive processing theory conceives the continuous evaluation of sensory uncertainty, influencing the impact of sensory prediction errors (Clark, 2016).

A test-retest approach confirmed that the observed behavioral performance metrics were reliable. Interestingly, there were important individual differences in how participants dealt with the difficulty and variability of the task, thus probably reflecting a ‘fingerprint’ of well-differentiated traits (Franklin & Wolpert, 2011; Trevino, Beltran-Navarro, et al., 2021; Tsutsui et al., 2019). Several executive functions (EF) are associated with complex cognitive tasks and adaptive motor control, especially those that involve changing conditions (Brosowsky & Egner, 2021). Cognitive aspects like (sustained) visual selective/spatial attention, stimulus detection, contrast sensitivity, visual search efficiency, switching/shifting (quick attentional adjustments, flexible allocation of attentional resources, cognitive flexibility, inhibitory control (cognitive inhibition, by not allowing the entry of stimuli that are not related to the task), visual short-term memory, working memory, decision making, and executive control of movement, could participate in our chasing/escaping task. Because our task emulates a complex problem, it could require the simultaneous participation of multiple cognitive processes. Also, the recruitment of EF seems to depend on task difficulty: more complex (less automated) tasks require these functions to a greater degree than easier (more automated) tasks (Diamond, 2013; Lezak et al., 2004).

The solution of the visuomotor task requires the interaction between perceptual and motor systems across many brain regions. The primary visual cortex (V1) first extracts information about the moving stimuli and sends it to the posterior parietal area to control visually-guided behavior. Parietal activity in mammals has been linked to visually guided real-time motor plan updates, perception of control over external stimuli, and learning task rules (Clancy & Mrsic-Flogel, 2021). The participation of sensory, motor, and attentional signals within the posterior parietal cortex allows shifting attention and response, a prerequisite for dealing with tasks that involve spatial processing (Lezak et al., 2004). Neurons in the dorsal anterior cingulate cortex (dACC) track position, velocity, and acceleration and have been implicated in prediction, prospection, and chasing moving objects (Yoo et al., 2020). In primates, the primary motor (M1) cortex encodes information about the interception and joystick’s movement (Merchant & Georgopoulos, 2006; Saxena et al., 2022). The cerebellum has also been considered a main region involved in impedance regulation (Franklin & Wolpert, 2011), and supports fine eye-hand coordination in visuomotor tasks (Miall et al., 2001). The lateral cerebellum receives sensory and motor inputs and mediates coordination control (Vercher et al., 2003). Some evidence also points to cerebellar participation in higher cognitive processes like language (Kawato, 1999).

Solving the visuomotor task combined visual and proprioceptive information. Indeed, on-line movement control demands the complex integration of peripheral sensory feed-back signals with predicted central feed-forward signals (Gritsenko et al., 2009). Here, the sense of movement control emerges with the capacity to perceive a causal relationship between internally generated actions and the outcomes in the external world (Clancy & Mrsic-Flogel, 2021). Both perceptual and action tasks require the capacity to follow moving objects visually. It is well known that when participants use their eyes to track a moving target, their visual scanning paths consist of smooth chase episodes interrupted by catch-up saccades when the location and/or velocity errors increase (Danion & Flanagan, 2018). Thus, it is essential to visually follow moving objects with fluid pursuit eye movements in both perceptual and visuomotor tasks. Since eye movements are often relatively short, it has been shown that the eyes tend to begin to move before the hand does. Therefore, the control of saccadic eye movements and goal-directed hand motions ends up being interdependent, as evidenced by the reliance of visuomotor pursuit on multimodal sensory streams to achieve target goals (Gritsenko et al., 2009; Hocherman & Levy, 2000; Niehorster et al., 2015; Vercher et al., 2003). Our task could be used in future studies to study self-monitoring (supervision of the execution of ongoing activities) and to assess the relative contributions of visual and non-visual inputs to visuomotor performance and the efficacy of different error detection and correction strategies. One interesting possibility is that participants, having acquired the models of the task’s dynamic properties, could exhibit the ability to anticipate collisions and strategically position their gaze accordingly. This anticipatory behavior would manifest in the form of “pro-active saccades”, where eye movements land in the anticipated collision location in advance. Such action control would rely on predictions rather than solely on perceptual information (Clark, 2016; Tatler et al., 2011).

Chasing strategies tend to become optimized over time (Furuichi, 2002). For instance, prediction accuracy increases when participants consider the statistical properties of the target’s motion. Thus, target interception includes determining the optimal speed and interception angle (Ghose et al., 2006; Warren, 2006). Our analytical approach allowed us to detect dynamic changes in pursuit *vs.* interception strategies during chasing trials. Escape behaviors also involved complicated sensorimotor control. Many animal species utilize escape reactions as their primary defense against predators. High accelerations, frequently coupled with a sudden change in direction, are typical escape responses to move the prey away from the attacker (Corcoran & Conner, 2016; Domenici et al., 2011), and the best escape angle depends on the speed difference between predator and prey (Corcoran & Conner, 2016). Accordingly, the relative velocity and inter-dot distance (IDD) are crucial metrics to determine escape interactions in sports (Tsutsui et al., 2019).

Chasing strategies in visuomotor coordination can be approached and controlled in various ways. One well-known strategy is classical pursuit, where the pursuer continuously aims directly at the target by moving toward its current location. This approach is commonly observed in animals and humans during prey chasing (Varennes et al., 2020). However, the delay in biological motor control is significant. For instance, the extraction of object velocity takes about 50–75 ms, and the latency for visual input on arm motions ranges between 150 and 250 milliseconds (Mrotek & Soechting, 2007). As a result, the time delays caused by brain transmission, muscle force production, and effector inertia render purely reactive behavior ineffective (Franklin & Wolpert, 2011; Wolpert et al., 1995). Reaching the target at the right position and time is crucial to intercept a moving target successfully. Since moving towards that position takes time, chasers must anticipate where to best intercept the target. Interception strategies involve predictive and/or prospective control (Zhao & Warren, 2015). Predictive control relies on pre-planned actions based on prior knowledge. It assumes familiarity with the target’s location or timing of contact, enabling accurate planning and execution. In this case, predictive control foresees that there should not be any late modifications to the planned movement (Franklin & Wolpert, 2011; Kawato, 1999; Kording & Wolpert, 2004; Panchuk & Vickers, 2009; Piray & Daw, 2020). Prospective strategies, on the other hand, continuously update movements to successfully intercept the target without requiring prior knowledge of the exact contact point (Bastin et al., 2006; Chai et al., 2019; Panchuk & Vickers, 2009; Varennes et al., 2020; Zhao & Warren, 2015). These interceptive actions rely on on-line sensory signals. Thus, while predictive control provides planned responses, prospective control focuses on the real-time regulation of movements based on a continuous stream of perceptual information. Both strategies have their advantages and can be utilized depending on the task requirements and the level of uncertainty in target motion (Zhao & Warren, 2015).

One of the main contributions of this work is the development of a method for identifying pursuit *vs.* interception behavior. We created a geometric model that probabilistically detects individual differences and dynamic changes in pursuit and interception behavior within chasing trials. Using this method, we discovered a significant increase in adopting a late interception strategy hundreds of milliseconds before chasers collided with the target. In this case, the variable *v_T_*, but not AR, played a crucial role in determining the percentage of the adopted late interception strategy (Fajen & Warren, 2004). One possibility is that this late interception approach gives decision-makers more time to gather information and formulate their chasing plan. Additionally, it supports the idea of on-line visually mediated adjustments of joystick motion during interception. From this perspective, it would be reasonable to assume that interception involves comparing anticipated and sensed target positions to assess a ‘prediction error’. Participants may utilize the results of such a comparison to command target interception (Mrotek & Soechting, 2007). Therefore, an on-line strategy could explain the delay involved in the late adoption of an interception strategy and the consistent performance observed with different AR and FDI variances. However, it is important to note that our current analysis cannot exclude the possibility of a hybrid control solution combining real-time visual information with an offline predictive model-based strategy (Zhao & Warren, 2015). Previous studies have utilized mathematical models to investigate the information sources involved in locomotor interception, specifically in tasks that require walking to intercept or escape a target that moves in depth from an embedded 1st-person viewpoint (Chardenon et al., 2005; Fajen & Warren, 2004, 2007). Our task differs in that it involves controlling a dot to intercept a moving target on a screen, adopting a 3rd-person viewpoint. Employing similar mathematical models to analyze our dataset could provide valuable insights into the potential interception strategies specific to our task context.

Our task allows to characterize chasing and escaping behavior in adults and children/youngsters. Neonates begin to acquire their capacity to grab moving objects soon after birth and, by 36 weeks of age, can appropriately intercept a moving item. Instead of viewing the moving object, newborns use a predictive strategy where the first movement is in the direction of the interception point. Children improve their reaching and gripping practices with age (Chinn et al., 2019). Furthermore, there is consensus that there is a parallel development in working and visuospatial memory (Alloway et al., 2006; Langan & Seidler, 2011), with a direct association with the development of inhibitory control (Sulik et al., 2018) and cognitive flexibility (Rigoli et al., 2012). However, little is known about how interception strategies change during development. As an initial step in this direction, our results showed how children increased their chasing/escaping performance and their interception strategies with age. We believe that the influence of task parameters and developmental stage on chasing and escaping performance is robust and should generalize to other similar tasks. Furthermore, different research assistants from our laboratories have obtained similar results in young students from different schools (unpublished observations). Therefore, we have no reason to believe that the results obtained depend on other characteristics of the participants, materials, or context.

Our study has several limitations that should be acknowledged. Firstly, the absence of eye movement recordings is a significant constraint. Incorporating an eye-tracking system into our task would greatly enhance our ability to investigate the visuomotor predictors associated with efficient interception. Indeed, the functional role of eye movements in guiding interception is not yet fully understood (Fooken et al., 2021), presenting opportunities for future experiments to explore this area. Another limitation is the lack of formal tests to explore the predominant interception strategies employed by participants. Specifically, we did not test whether the observed strategies involved predictive control, relying on prior knowledge and pre-planned actions, or prospective control, which entails real-time adjustments based on continuous perceptual information. Performing formal tests in this regard would lead to a more precise characterization of the interception strategies employed and result in a clearer comprehension of the underlying cognitive processes involved in target interception. An important limitation of our interception trajectory detection method stems from the relatively short distances covered by the target during each FDI. As a result, there is an overlap in the mean value of the error distributions for both pursuit and interception strategies when the user is far away from the target. This overlap presents a challenge in distinguishing between the two strategies at longer distances. However, our observations indicate that this convergence does not occur at shorter distances, allowing us to differentiate between the two strategies using the %OIA metric. Another limitation in our task was the constant speed of the target, regardless of the amplitude of the directional change imposed on it. However, it is widely known that biological motion typically adheres to the 2/3 power law (Viviani & Flash, 1995). This law states that as curvature increases, movement tends to slow down, and this relationship can be attributed to both mechanical and neural factors. Therefore, it would be valuable to investigate how performance is affected when the target’s speed follows the 2/3 power law while changing its movement direction, thus providing insights into the potential improvement in performance under more ecologically realistic conditions.

The interactive nature of our task, designed as a game, adds an element of excitement and motivation, making it suitable for engaging younger participants. Examining interception strategies during infancy is particularly important because our methods enable a close examination of the factors involved in the development of this cognitive skill. Our approach provides highly detailed metrics for characterizing the interception of moving targets, offering potential means to quantify the proper development of internal predictive and prospective strategies and motor skills (Franklin et al., 2008). On the other end of the spectrum, our task holds the potential for monitoring cognitive decline in older participants and studying the progression of psychiatric disorders. It can also be employed to investigate how prediction errors influence learning and decision-making, and how individuals cope with visuomotor discrepancies (Gidley Larson et al., 2008). In practical scenarios, such as rehabilitation, gaining insights into how we acquire knowledge from our mistakes is crucial for understanding how the brain forms representations of prior expectations and probabilities in a Bayesian manner (Franklin & Wolpert, 2011; Sedaghat-Nejad & Shadmehr, 2021). Furthermore, our task and methods present a promising potential for diagnostic purposes, as they allow the independent study of the visual and motor mechanisms implicated in continuously modifying and updating our responses.

### Constraints on Generality

All adult participants in our study were enrolled at Universidad de Guadalajara, Mexico’s second-largest public college. The child participants, on the other hand, were matriculated at Instituto de Ciencias, the largest private school in Guadalajara Jalisco, Mexico. It is worth noting that Guadalajara Jalisco is the second largest city in Mexico and attracts students from diverse socioeconomic backgrounds to both educational institutions. As a result, our sample includes participants from various socioeconomic levels, allowing for reasonable generalization of the results within this context. While our study focused on participants from these specific institutions and geographic locations, further research is needed to determine how our findings can be generalized to other populations and settings. Future studies involving participants from different educational backgrounds and geographic locations could contribute to a more comprehensive understanding of the generality of our results. By acknowledging these constraints on generality, we promote transparency and facilitate the interpretation and replication of our study in diverse contexts.

## Author note

The ideas and data presented in this manuscript have not been previously disseminated or shared. The content of this manuscript has not been presented at any conference or meeting, nor has it been posted on any listserv, shared on a website, or made available through any other platform prior to its submission to this journal.

## Availability of Data and Materials

Datasets from our experiments are available from Open Science Framework (OSF, https://osf.io/nsbhr).

## Author Contributions

Conceptualization: MT; Data curation: MT; Formal analysis: ST, MT; Funding acquisition: MT; Investigation: JV, BB, ST; Methodology: BB, ST, RM, MT; Project administration: MT; Resources: MT; Software: RM, MT; Supervision: BB, MT; Validation: BB, JV, MT; Visualization: ST, MT; Writing—original draft: MT; Writing—review and editing: RM, ST, BB, MT.

## Acknowledgments

We thank B. De la Torre-Valdovinos and P. Osuna Carrasco for sharing their lab space to run our experiments and for helping us to get participants for the study; H. Hinojiante and R. Torres for conducting some tests. Thanks to Dr. M. Torres Morán and her team from Coordinación de Investigación CUCBA, UdeG, for their help and support. Special thanks to the reviewers for helping us to improve our manuscript.

## References

Alloway, T. P., Gathercole, S. E., & Pickering, S. J. (2006). Verbal and visuospatial short-term and working memory in children: are they separable? Child Dev, 77(6), 1698–1716. https://doi.org/10.1111/j.1467-8624.2006.00968.x

Bastin, J., Craig, C., & Montagne, G. (2006). Prospective strategies underlie the control of interceptive actions. Hum Mov Sci, 25(6), 718–732. https://doi.org/10.1016/j.humov.2006.04.001

Behrens, T. E., Woolrich, M. W., Walton, M. E., & Rushworth, M. F. (2007). Learning the value of information in an uncertain world. Nat Neurosci, 10(9), 1214–1221. https://doi.org/10.1038/nn1954

Berens, P. (2009). CircStat: A MATLAB Toolbox for Circular Statistics. Journal of Statistical Software, 31(10), 1–21. https://doi.org/10.18637/jss.v031.i10

Brenner, E., & Smeets, J. B. (2015). How people achieve their amazing temporal precision in interception. J Vis, 15(3). https://doi.org/10.1167/15.3.8

Brosowsky, N. P., & Egner, T. (2021). Appealing to the cognitive miser: Using demand avoidance to modulate cognitive flexibility in cued and voluntary task switching. J Exp Psychol Hum Percept Perform, 47(10), 1329–1347. https://doi.org/10.1037/xhp0000942

Chai, R., Savvaris, A., & Chai, S. (2019). Integrated missile guidance and control using optimization-based predictive control. Nonlinear Dynamics, 96(2), 997–1015. https://doi.org/10.1007/s11071-019-04835-8

Chardenon, A., Montagne, G., Laurent, M., & Bootsma, R. J. (2005). A robust solution for dealing with environmental changes in intercepting moving balls. J Mot Behav, 37(1), 52–64. https://doi.org/10.3200/JMBR.37.1.52-62

Chinn, L. K., Noonan, C. F., Hoffmann, M., & Lockman, J. J. (2019). Development of Infant Reaching Strategies to Tactile Targets on the Face. Front Psychol, 10, 9. https://doi.org/10.3389/fpsyg.2019.00009

Clancy, K. B., & Mrsic-Flogel, T. D. (2021). The sensory representation of causally controlled objects. Neuron, 109(4), 677–689 e674. https://doi.org/10.1016/j.neuron.2020.12.001

Clark, A. (2016). Surfing Uncertainty: Prediction, Action, and the Embodied Mind. Oxford University Press. https://doi.org/10.1093/acprof:oso/9780190217013.001.0001

Corcoran, A. J., & Conner, W. E. (2016). How moths escape bats: predicting outcomes of predator-prey interactions. J Exp Biol, 219(Pt 17), 2704–2715. https://doi.org/10.1242/jeb.137638

Coudiere, A., Fernandez, E., de Rugy, A., & Danion, F. R. (2022). Asymmetrical transfer of adaptation between reaching and tracking: implications for feedforward and feedback processes. J Neurophysiol, 128(3), 480–493. https://doi.org/10.1152/jn.00547.2021

Danion, F. R., & Flanagan, J. R. (2018). Different gaze strategies during eye versus hand tracking of a moving target. Sci Rep, 8(1), 10059. https://doi.org/10.1038/s41598-018-28434-6

Dessing, J. C., Peper, C. L., Bullock, D., & Beek, P. J. (2005). How position, velocity, and temporal information combine in the prospective control of catching: data and model. J Cogn Neurosci, 17(4), 668–686. https://doi.org/10.1162/0898929053467604

Diamond, A. (2013). Executive functions. Annu Rev Psychol, 64, 135–168. https://doi.org/10.1146/annurev-psych-113011-143750

Domenici, P., Blagburn, J. M., & Bacon, J. P. (2011). Animal escapology II: escape trajectory case studies. J Exp Biol, 214(Pt 15), 2474–2494. https://doi.org/10.1242/jeb.053801

Ernst, M. O., & Banks, M. S. (2002). Humans integrate visual and haptic information in a statistically optimal fashion. Nature, 415(6870), 429–433. https://doi.org/10.1038/415429a

Fajen, B. R., & Warren, W. H. (2004). Visual guidance of intercepting a moving target on foot. Perception, 33(6), 689–715. https://doi.org/10.1068/p5236

Fajen, B. R., & Warren, W. H. (2007). Behavioral dynamics of intercepting a moving target. Exp Brain Res, 180(2), 303–319. https://doi.org/10.1007/s00221-007-0859-6

Faul, F., Erdfelder, E., Buchner, A., & Lang, A. G. (2009). Statistical power analyses using G*Power 3.1: tests for correlation and regression analyses. Behav Res Methods, 41(4), 1149–1160. https://doi.org/10.3758/BRM.41.4.1149

Festa-Bianchet, M., & Mysterud, A. (2018). Hunting and evolution: theory, evidence, and unknowns. Journal of Mammalogy, 99(6), 1281–1292. https://doi.org/10.1093/jmammal/gyy138

Fooken, J., Kreyenmeier, P., & Spering, M. (2021). The role of eye movements in manual interception: A mini-review. Vision Res, 183, 81–90. https://doi.org/10.1016/j.visres.2021.02.007

Franklin, D. W., Burdet, E., Tee, K. P., Osu, R., Chew, C. M., Milner, T. E., & Kawato, M. (2008). CNS learns stable, accurate, and efficient movements using a simple algorithm. J Neurosci, 28(44), 11165–11173. https://doi.org/10.1523/JNEUROSCI.3099-08.2008

Franklin, D. W., & Wolpert, D. M. (2011). Computational mechanisms of sensorimotor control. Neuron, 72(3), 425–442. https://doi.org/10.1016/j.neuron.2011.10.006

Furuichi, N. (2002). Dynamics between a predator and a prey switching two kinds of escape motions. J Theor Biol, 217(2), 159–166. https://doi.org/10.1006/jtbi.2002.3027

Ghose, K., Horiuchi, T. K., Krishnaprasad, P. S., & Moss, C. F. (2006). Echolocating bats use a nearly time-optimal strategy to intercept prey. PLoS Biol, 4(5), e108. https://doi.org/10.1371/journal.pbio.0040108

Gidley Larson, J. C., Bastian, A. J., Donchin, O., Shadmehr, R., & Mostofsky, S. H. (2008). Acquisition of internal models of motor tasks in children with autism. Brain, 131(Pt 11), 2894–2903. https://doi.org/10.1093/brain/awn226

Gritsenko, V., Yakovenko, S., & Kalaska, J. F. (2009). Integration of predictive feedforward and sensory feedback signals for online control of visually guided movement. J Neurophysiol, 102(2), 914–930. https://doi.org/10.1152/jn.91324.2008

Hocherman, S., & Levy, H. (2000). The role of feedback in manual tracking of visual targets. Percept Mot Skills, 90(3 Pt 2), 1235-1248. https://doi.org/10.2466/pms.2000.90.3c.1235

Howland, H. C. (1974). Optimal strategies for predator avoidance: the relative importance of speed and manoeuvrability. J Theor Biol, 47(2), 333–350. https://doi.org/10.1016/0022-5193(74)90202-1

Kawato, M. (1999). Internal models for motor control and trajectory planning. Curr Opin Neurobiol, 9(6), 718–727. https://doi.org/10.1016/s0959-4388(99)00028-8

Knill, D. C., & Pouget, A. (2004). The Bayesian brain: the role of uncertainty in neural coding and computation. Trends Neurosci, 27(12), 712–719. https://doi.org/10.1016/j.tins.2004.10.007

Kording, K. P., & Wolpert, D. M. (2004). Bayesian integration in sensorimotor learning. Nature, 427(6971), 244–247. https://doi.org/10.1038/nature02169

Kwon, O. S., & Knill, D. C. (2013). The brain uses adaptive internal models of scene statistics for sensorimotor estimation and planning. Proc Natl Acad Sci U S A, 110(11), E1064–1073. https://doi.org/10.1073/pnas.1214869110

Lam, S. Y., & Zenon, A. (2021). Information Rate in Humans during Visuomotor Tracking. Entropy (Basel), 23(2). https://doi.org/10.3390/e23020228

Langan, J., & Seidler, R. D. (2011). Age differences in spatial working memory contributions to visuomotor adaptation and transfer. Behav Brain Res, 225(1), 160–168. https://doi.org/10.1016/j.bbr.2011.07.014

Lenroot, R. K., & Giedd, J. N. (2006). Brain development in children and adolescents: insights from anatomical magnetic resonance imaging. Neurosci Biobehav Rev, 30(6), 718–729. https://doi.org/10.1016/j.neubiorev.2006.06.001

Lezak, M. D., Howieson, D. B., Loring, D. W., & Fischer, J. S. (2004). Neuropsychological Assessment (4th ed.). Oxford University Press.

Mehta, B., & Schaal, S. (2002). Forward models in visuomotor control. J Neurophysiol, 88(2), 942–953. https://doi.org/10.1152/jn.2002.88.2.942

Merchant, H., & Georgopoulos, A. P. (2006). Neurophysiology of perceptual and motor aspects of interception. J Neurophysiol, 95(1), 1–13. https://doi.org/10.1152/jn.00422.2005

Miall, R. C., Reckess, G. Z., & Imamizu, H. (2001). The cerebellum coordinates eye and hand tracking movements. Nat Neurosci, 4(6), 638–644. https://doi.org/10.1038/88465

Mischiati, M., Lin, H. T., Herold, P., Imler, E., Olberg, R., & Leonardo, A. (2015). Internal models direct dragonfly interception steering. Nature, 517(7534), 333–338. https://doi.org/10.1038/nature14045

Mrotek, L. A., & Soechting, J. F. (2007). Target interception: hand-eye coordination and strategies. J Neurosci, 27(27), 7297–7309. https://doi.org/10.1523/JNEUROSCI.2046-07.2007

Niehorster, D. C., Siu, W. W., & Li, L. (2015). Manual tracking enhances smooth pursuit eye movements. J Vis, 15(15), 11. https://doi.org/10.1167/15.15.11

Panchuk, D., & Vickers, J. N. (2009). Using spatial occlusion to explore the control strategies used in rapid interceptive actions: Predictive or prospective control? J Sports Sci, 27(12), 1249–1260. https://doi.org/10.1080/02640410903156449

Piray, P., & Daw, N. D. (2020). A simple model for learning in volatile environments. PLoS Comput Biol, 16(7), e1007963. https://doi.org/10.1371/journal.pcbi.1007963

Rigoli, D., Piek, J. P., Kane, R., & Oosterlaan, J. (2012). An examination of the relationship between motor coordination and executive functions in adolescents. Dev Med Child Neurol, 54(11), 1025–1031. https://doi.org/10.1111/j.1469-8749.2012.04403.x

Saxena, S., Russo, A. A., Cunningham, J., & Churchland, M. M. (2022). Motor cortex activity across movement speeds is predicted by network-level strategies for generating muscle activity. Elife, 11. https://doi.org/10.7554/eLife.67620

Schroeger, A., Tolentino-Castro, J. W., Raab, M., & Canal-Bruland, R. (2021). Effects of visual blur and contrast on spatial and temporal precision in manual interception. Exp Brain Res, 239(11), 3343–3358. https://doi.org/10.1007/s00221-021-06184-8

Sedaghat-Nejad, E., & Shadmehr, R. (2021). The cost of correcting for error during sensorimotor adaptation. Proc Natl Acad Sci U S A, 118(40). https://doi.org/10.1073/pnas.2101717118

Soechting, J. F., Juveli, J. Z., & Rao, H. M. (2009). Models for the extrapolation of target motion for manual interception. J Neurophysiol, 102(3), 1491–1502. https://doi.org/10.1152/jn.00398.2009

Sulik, M. J., Haft, S. L., & Obradović, J. (2018). Visual-motor integration, executive functions, and academic achievement: Concurrent and longitudinal relations in late elementary school.. Early education and development, 29(7), 956–970. https://doi.org/https://doi.org/10.1080/10409289.2018.1442097

Tatler, B. W., Hayhoe, M. M., Land, M. F., & Ballard, D. H. (2011). Eye guidance in natural vision: reinterpreting salience. J Vis, 11(5), 5. https://doi.org/10.1167/11.5.5

Todorov, E., & Jordan, M. I. (2002). Optimal feedback control as a theory of motor coordination. Nat Neurosci, 5(11), 1226–1235. https://doi.org/10.1038/nn963

Trevino, M., Beltran-Navarro, B., Leon, R. M. Y., & Matute, E. (2021). Clustering of neuropsychological traits of preschoolers. Sci Rep, 11(1), 6533. https://doi.org/10.1038/s41598-021-85891-2

Trevino, M., Castiello, S., Arias-Carrion, O., De la Torre-Valdovinos, B., & Coss, Y. L. R. M. (2021). Isomorphic decisional biases across perceptual tasks. PLoS One, 16(1), e0245890. https://doi.org/10.1371/journal.pone.0245890

Trevino, M., Medina-Coss, Y. L. R., & Haro, B. (2020). Adaptive Choice Biases in Mice and Humans. Front Behav Neurosci, 14, 99. https://doi.org/10.3389/fnbeh.2020.00099

Tsutsui, K., Shinya, M., & Kudo, K. (2019). Underlying structure in the dynamics of chase and escape interactions. Sci Rep, 9(1), 15051. https://doi.org/10.1038/s41598-019-51524-y

Tsutsui, K., Shinya, M., & Kudo, K. (2020). Human Navigational Strategy for Intercepting an Erratically Moving Target in Chase and Escape Interactions. J Mot Behav, 52(6), 750–760. https://doi.org/10.1080/00222895.2019.1692331

Varennes, L., Krapp, H. G., & Viollet, S. (2020). Two pursuit strategies for a single sensorimotor control task in blowfly. Sci Rep, 10(1), 20762. https://doi.org/10.1038/s41598-020-77607-9

Vercher, J. L., Sares, F., Blouin, J., Bourdin, C., & Gauthier, G. (2003). Role of sensory information in updating internal models of the effector during arm tracking. Prog Brain Res, 142, 203–222. https://doi.org/10.1016/S0079-6123(03)42015-3

Viviani, P., & Flash, T. (1995). Minimum-jerk, two-thirds power law, and isochrony: converging approaches to movement planning. J Exp Psychol Hum Percept Perform, 21(1), 32–53. https://doi.org/10.1037//0096-1523.21.1.32

Warren, W. H. (2006). The dynamics of perception and action. Psychol Rev, 113(2), 358–389. https://doi.org/10.1037/0033-295X.113.2.358

Wolpert, D. M., Ghahramani, Z., & Jordan, M. I. (1995). An internal model for sensorimotor integration. Science, 269(5232), 1880–1882. https://doi.org/10.1126/science.7569931

Yoo, S. B. M., Tu, J. C., Piantadosi, S. T., & Hayden, B. Y. (2020). The neural basis of predictive pursuit. Nat Neurosci, 23(2), 252–259. https://doi.org/10.1038/s41593-019-0561-6

Zago, M., McIntyre, J., Senot, P., & Lacquaniti, F. (2009). Visuo-motor coordination and internal models for object interception. Exp Brain Res, 192(4), 571–604. https://doi.org/10.1007/s00221-008-1691-3

Zhao, H., & Warren, W. H. (2015). On-line and model-based approaches to the visual control of action. Vision Res, 110(Pt B), 190–202. https://doi.org/10.1016/j.visres.2014.10.008

