## Supplementary Material for "Directional uncertainty in chase and escape dynamics"

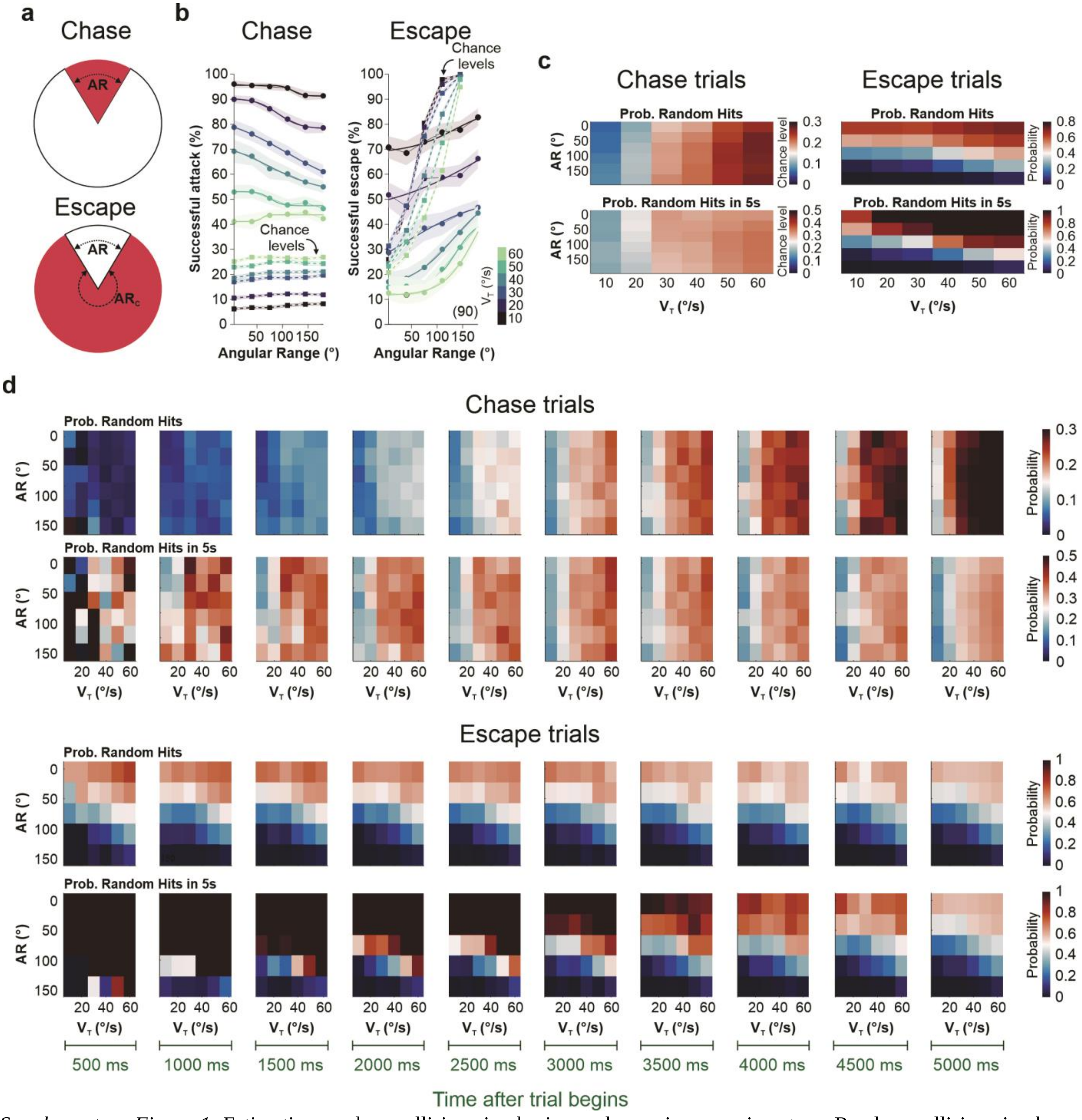


*Supplementary Figure 1*. Estimating random collisions in chasing and escaping experiments. **a** Random collisions in chase trials were estimated by simulating alternative computer trajectories using the same $v_T$ and AR as those used for the original trials (upper panel). For escape trials, we calculated random collisions by using the same $v_T$ and $AR_C$ (the complement of AR; lower panel; see **Methods**). **b** Psychometric curves from groups of participants solving the chase (right) and escape trials (left). Squares represent chance levels obtained by calculating random collisions. **c** Color panels show the probability of random collision during the analyzed trials (*i.e.*, using paths from the actual traces; upper panels), and the probability of random collision in a time window of 5 s (equivalent to max. trial duration). Both probability colormaps are shown as a function of AR and $v_T$ with an FDI = 100 ms. **d** Similar analyses as before but breaking them down into cumulative 500 ms windows, from the beginning of the trial to the max duration of a trial. Labels for the corresponding time windows on the bottom. Number of participants in parentheses.

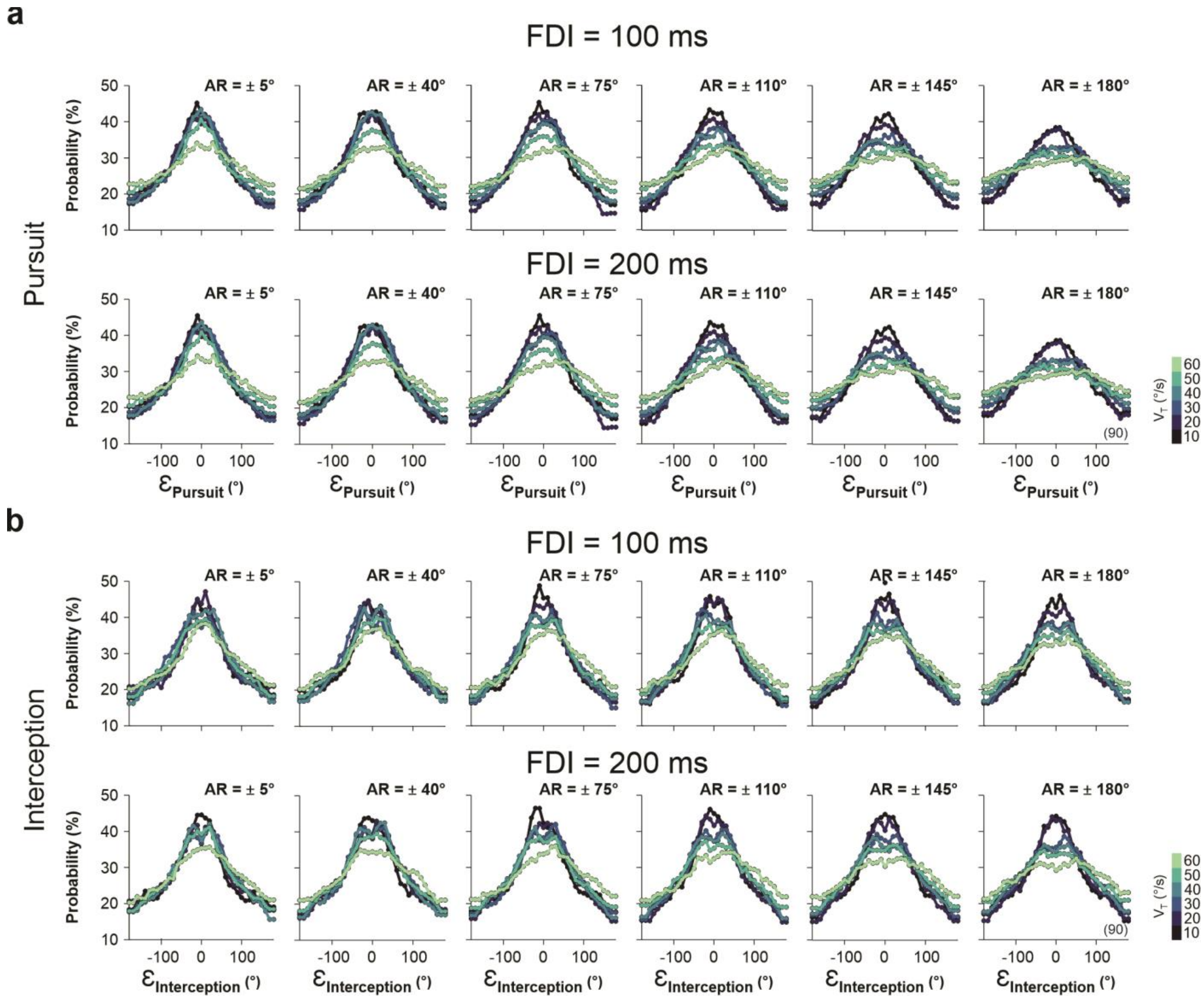


*Supplementary Figure 2*. Gaussian fits to tracking and interception distributions of angular errors. This figure is complementary to *Figure 4b*. The panels show the Gaussian fits to the observed distributions of angular errors for pursuit (**a**) and interception (**b**) errors (see **Methods**). Colorbar represents $v_T$. Number of participants per experiment in parentheses.

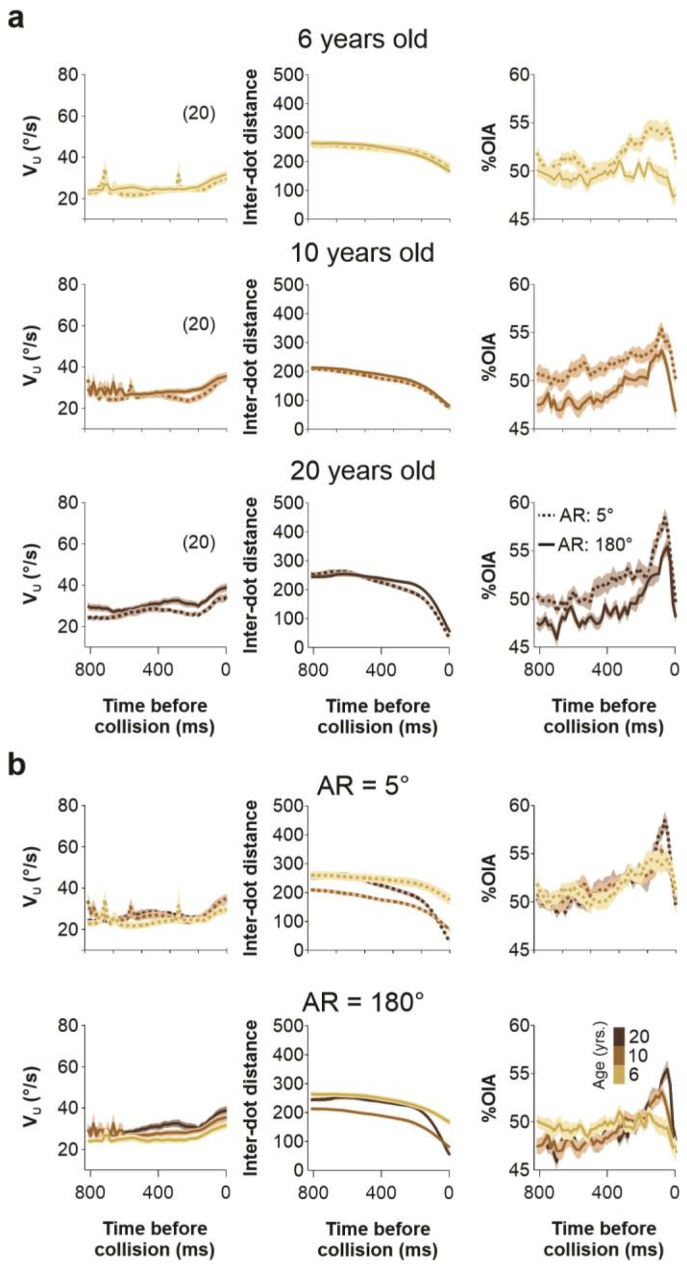


*Supplementary Figure 3*. Changes in chasing/escaping $v_U$ and IDD through development. This figure is complementary to *Figure 6c,d*. **a** Collision triggered averages of user speed ($v_U$), inter-dot distance (IDD), and %OIA (see **Method**) before colliding with the target from infants and youngsters belonging to three groups: 5, 10 and 20 years of age. **b** Same analysis but with over imposed traces for the different age groups, which are represented in the colorbar to the right. Number of participants in parentheses.

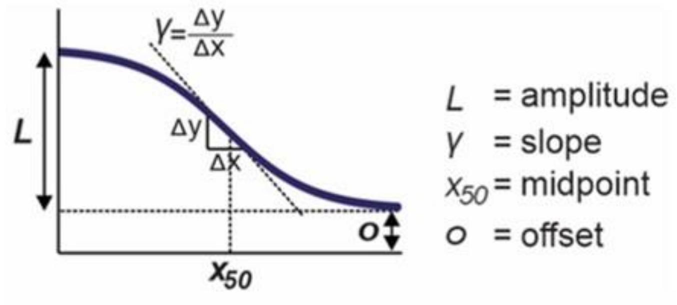


**Chase**

| | FDI = 100 ms | | | | FDI = 200 ms | | | | |
|---|---|---|---|---|---|---|---|---|---|
| $V_{CT}$ | $L$ | γ | $x_{50}$ | o | $L$ | γ | $x_{50}$ | o | n |
| 10 | 0.13 ± 0.06 | 0.45 ± 0.10 | 67.81 ± 8.394 | 0.87 ± 0.03 | 0.12 ± 0.06 | 0.44 ± 0.11 | 94.89 ± 15.90 | 0.92 ± 0.03 | 15 |
| 20 | 0.21 ± 0.05 | 0.27 ± 0.09 | 97.74 ± 16.46 | 0.76 ± 0.02 | 0.24 ± 0.07 | 0.39 ± 0.10 | 90.69 ± 16.24 | 0.78 ± 0.03 | 15 |
| 30 | 0.19 ± 0.02 | 0.35 ± 0.09 | 79.94 ± 9.72 | 0.60 ± 0.02 | 0.15 ± 0.03 | 0.52 ± 0.11 | 73.96 ± 9.35 | 0.64 ± 0.03 | 15 |
| 40 | 0.30 ± 0.09 | 0.49 ± 0.10 | 108.91 ± 13.96 | 0.51 ± 0.03 | 0.31 ± 0.09 | 0.42 ± 0.10 | 97.39 ± 14.14 | 0.52 ± 0.05 | 15 |
| 50 | 0.20 ± 0.06 | 0.46 ± 0.10 | 104.75 ± 14.84 | 0.41 ± 0.04 | 0.23 ± 0.07 | 0.53 ± 0.08 | 109.70 ± 17.38 | 0.47 ± 0.03 | 15 |
| 60 | 0.16 ± 0.06 | 0.51 ± 0.11 | 93.01 ± 14.96 | 0.35 ± 0.03 | 0.09 ± 0.02 | 0.69 ± 0.09 | 99.74 ± 11.49 | 0.39 ± 0.01 | 15 |

**Escape**

| | FDI = 100 ms | | | | FDI = 200 ms | | | | |
|---|---|---|---|---|---|---|---|---|---|
| $V_{CT}$ | $L$ | γ | $x_{50}$ | o | $L$ | γ | $x_{50}$ | o | n |
| 10 | 0.31 ± 0.07 | 0.59 ± 0.09 | 108.48 ± 13.22 | 0.65 ± 0.05 | 0.33 ± 0.07 | 0.46 ± 0.09 | 122.25 ± 12.37 | 0.51 ± 0.03 | 15 |
| 20 | 0.31 ± 0.07 | 0.60 ± 0.09 | 99.19 ± 11.63 | 0.45 ± 0.04 | 0.37 ± 0.06 | 0.45 ± 0.07 | 124.70 ± 12.55 | 0.32 ± 0.02 | 15 |
| 30 | 0.29 ± 0.06 | 0.60 ± 0.07 | 96.87 ± 10.72 | 0.27 ± 0.04 | 0.44 ± 0.06 | 0.50 ± 0.08 | 119.17 ± 11.42 | 0.21 ± 0.03 | 15 |
| 40 | 0.42 ± 0.07 | 0.41 ± 0.08 | 124.56 ± 10.58 | 0.16 ± 0.03 | 0.45 ± 0.06 | 0.33 ± 0.08 | 125.48 ± 9.21 | 0.17 ± 0.03 | 15 |
| 50 | 0.45 ± 0.08 | 0.39 ± 0.08 | 143.88 ± 9.26 | 0.13 ± 0.03 | 0.47 ± 0.06 | 0.40 ± 0.07 | 119.93 ± 12.05 | 0.14 ± 0.03 | 15 |
| 60 | 0.35 ± 0.06 | 0.41 ± 0.09 | 143.38 ± 9.73 | 0.11 ± 0.03 | 0.53 ± 0.06 | 0.28 ± 0.08 | 115.71 ± 10.22 | 0.12 ± 0.03 | 15 |

*Supplementary Table 1*. Parameter fits from the psychometric curves from chasing and escaping experiments. This table shows averages ± S.E.M.

**Chase**

| $V_{CT}$ | FDI = 100 ms | | | | FDI = 200 ms | | | | |
|---|---|---|---|---|---|---|---|---|---|
| | $L$ | γ | $x_{50}$ | o | $L$ | γ | $x_{50}$ | o | n |
| 10 | 0.05 ± 0.01 | 0.46 | 74.70 ± 6.80 | 0.91 ± 0.01 | 0.05 ± 0.02 | 0.46 | 79.67 ± 12.74 | 0.94 ± 0.01 | 15 |
| 20 | 0.11 ± 0.02 | 0.46 | 80.08 ± 6.24 | 0.79 ± 0.02 | 0.09 ± 0.01 | 0.46 | 77.03 ± 7.80 | 0.84 ± 0.01 | 15 |
| 30 | 0.15 ± 0.01 | 0.46 | 89.42 ± 6.78 | 0.63 ± 0.02 | 0.11 ± 0.02 | 0.46 | 78.48 ± 8.61 | 0.67 ± 0.03 | 15 |
| 40 | 0.17 ± 0.03 | 0.46 | 95.83 ± 10.54 | 0.55 ± 0.02 | 0.16 ± 0.03 | 0.46 | 98.80 ± 11.42 | 0.58 ± 0.04 | 15 |
| 50 | 0.08 ± 0.02 | 0.46 | 86.41 ± 10.48 | 0.46 ± 0.02 | 0.15 ± 0.04 | 0.46 | 88.62 ± 12.91 | 0.47 ± 0.03 | 15 |
| 60 | 0.14 ± 0.05 | 0.46 | 79.57 ± 10.73 | 0.38 ± 0.02 | 0.08 ± 0.02 | 0.46 | 87.93 ± 10.22 | 0.39 ± 0.01 | 15 |

**Escape**

| $V_{CT}$ | FDI = 100 ms | | | | FDI = 200 ms | | | | |
|---|---|---|---|---|---|---|---|---|---|
| | $L$ | γ | $x_{50}$ | o | $L$ | γ | $x_{50}$ | o | n |
| 10 | 0.14 ± 0.02 | 0.46 | 89.95 ± 8.72 | 0.68 ± 0.04 | 0.17 ± 0.02 | 0.46 | 80.42 ± 7.82 | 0.50 ± 0.03 | 15 |
| 20 | 0.18 ± 0.03 | 0.46 | 82.68 ± 8.42 | 0.47 ± 0.04 | 0.26 ± 0.04 | 0.46 | 109.27 ± 8.05 | 0.32 ± 0.02 | 15 |
| 30 | 0.23 ± 0.02 | 0.46 | 88.28 ± 9.50 | 0.28 ± 0.04 | 0.26 ± 0.02 | 0.46 | 92.41 ± 7.74 | 0.24 ± 0.03 | 15 |
| 40 | 0.25 ± 0.02 | 0.46 | 99.43 ± 7.65 | 0.17 ± 0.03 | 0.31 ± 0.03 | 0.46 | 114.24 ± 7.12 | 0.20 ± 0.03 | 15 |
| 50 | 0.26 ± 0.02 | 0.46 | 124.20 ± 5.62 | 0.14 ± 0.03 | 0.30 ± 0.02 | 0.46 | 101.68 ± 8.14 | 0.17 ± 0.03 | 15 |
| 60 | 0.29 ± 0.05 | 0.46 | 125.69 ± 7.54 | 0.11 ± 0.03 | 0.34 ± 0.02 | 0.46 | 97.22 ± 6.41 | 0.15 ± 0.03 | 15 |

*Supplementary Table 2.* Parameter fits from the psychometric curves from chasing and escaping experiments with a fixed γ for all groups. This table shows averages ± S.E.M.

**Chase**

| | FDI = 100 ms | | | | FDI = 200 ms | | | | |
|---|---|---|---|---|---|---|---|---|---|
| $V_{CT}$ | *L* | γ | $x_{50}$ | o | *L* | γ | $x_{50}$ | o | n |
| 10 | 0.37 ± 0.12 | 0.29 ± 0.12 | 93.74 | 0.76 ± 0.06 | 0.33 ± 0.11 | 0.34 ± 0.13 | 93.74 | 0.79 ± 0.05 | 15 |
| 20 | 0.37 ± 0.11 | 0.20 ± 0.10 | 93.74 | 0.66 ± 0.05 | 0.52 ± 0.12 | 0.28 ± 0.12 | 93.74 | 0.63 ± 0.06 | 15 |
| 30 | 0.38 ± 0.11 | 0.29 ± 0.12 | 93.74 | 0.52 ± 0.06 | 0.40 ± 0.12 | 0.24 ± 0.11 | 93.74 | 0.53 ± 0.06 | 15 |
| 40 | 0.48 ± 0.12 | 0.23 ± 0.11 | 93.74 | 0.39 ± 0.06 | 0.45 ± 0.11 | 0.16 ± 0.09 | 93.74 | 0.42 ± 0.06 | 15 |
| 50 | 0.36 ± 0.10 | 0.22 ± 0.11 | 93.74 | 0.32 ± 0.05 | 0.57 ± 0.11 | 0.15 ± 0.09 | 93.74 | 0.24 ± 0.06 | 15 |
| 60 | 0.37 ± 0.10 | 0.10 ± 0.07 | 93.74 | 0.24 ± 0.05 | 0.18 ± 0.08 | 0.32 ± 0.12 | 93.74 | 0.34 ± 0.04 | 15 |

**Escape**

| | FDI = 100 ms | | | | FDI = 200 ms | | | | |
|---|---|---|---|---|---|---|---|---|---|
| $V_{CT}$ | *L* | γ | $x_{50}$ | o | *L* | γ | $x_{50}$ | o | n |
| 10 | 0.29 ± 0.09 | 0.20 ± 0.08 | 93.74 | 0.60 ± 0.05 | 0.33 ± 0.11 | 0.34 ± 0.13 | 93.74 | 0.79 ± 0.05 | 15 |
| 20 | 0.50 ± 0.09 | 0.23 ± 0.09 | 93.74 | 0.32 ± 0.05 | 0.52 ± 0.12 | 0.28 ± 0.12 | 93.74 | 0.63 ± 0.06 | 15 |
| 30 | 0.47 ± 0.07 | 0.18 ± 0.08 | 93.74 | 0.15 ± 0.04 | 0.40 ± 0.12 | 0.24 ± 0.11 | 93.74 | 0.53 ± 0.06 | 15 |
| 40 | 0.41 ± 0.06 | 0.18 ± 0.08 | 93.74 | 0.08 ± 0.02 | 0.45 ± 0.11 | 0.16 ± 0.09 | 93.74 | 0.42 ± 0.06 | 15 |
| 50 | 0.27 ± 0.03 | 0.18 ± 0.07 | 93.74 | 0.10 ± 0.03 | 0.57 ± 0.11 | 0.15 ± 0.09 | 93.74 | 0.24 ± 0.06 | 15 |
| 60 | 0.24 ± 0.04 | 0.19 ± 0.08 | 93.74 | 0.08 ± 0.03 | 0.18 ± 0.08 | 0.32 ± 0.12 | 93.74 | 0.34 ± 0.04 | 15 |

*Supplementary Table 3.* Parameter fits from the psychometric curves from chasing and escaping experiments with a fixed $x_{50}$ for all groups. This table shows averages ± S.E.M.
